# Engineering Material Properties of Transcription Factor Condensates to Control Gene Expression in Mammalian Cells and Mice

**DOI:** 10.1101/2023.10.16.562453

**Authors:** Alexandra A.M. Fischer, Hanah B. Robertson, Deqiang Kong, Merlin M. Grimm, Jakob Grether, Johanna Groth, Carsten Baltes, Manfred Fliegauf, Franziska Lautenschläger, Bodo Grimbacher, Haifeng Ye, Volkhard Helms, Wilfried Weber

**Affiliations:** Signalling Research Centres BIOSS and CIBSS, University of Freiburg, Schänzlestraße 18, 79104 Freiburg, Germany; Faculty of Biology, University of Freiburg, Schänzlestraße 1, 79104 Freiburg, Germany; Spemann Graduate School of Biology and Medicine (SGBM), University of Freiburg, Albertstraße 21a, 79104 Freiburg, Germany; INM – Leibniz Institute for New Materials, Campus D2 2, 66123 Saarbrücken, Germany; Saarland University, Department of Materials Science and Engineering, Campus D2 2, 66123 Saarbrücken, Germany; Synthetic Biology and Biomedical Engineering Laboratory, Biomedical Synthetic Biology Research Center, Shanghai Key Laboratory of Regulatory Biology, Institute of Biomedical Sciences and School of Life Sciences East China Normal University, Dongchuan Road 500, Shanghai 200241, China; Biberach University of Applied Sciences, Karlstraße 6-11, 88400 Biberach an der Riß, Germany; Institute for Immunodeficiency, Center for Chronic Immunodeficiency (CCI) Medical Center, Faculty of Medicine, University of Freiburg, Breisacherstr. 115, 79106 Freiburg, Germany; DZIF – German Center for Infection Research, Deutsches Zentrum für Infektionsforschung e.V., Inhoffenstr. 7, 38124 Braunschweig, Germany; RESIST – Cluster of Excellence 2155 to Hanover Medical School, Medizinische Hochschule Hannover, Carl-Neuberg-Str. 1, 30625 Hannover, Germany; CIBSS – Centre for Integrative Biological Signalling Studies, University of Freiburg, Schänzlestraße 18, 79104 Freiburg, Germany; Saarland University, Center for Bioinformatics, Saarland Informatics Campus, 66123 Saarbrücken, Germany; Saarland University, Department of Experimental Physics and Center for Biophysics, 66123 Saarbrücken, Germany

**Keywords:** biomolecular condensates, liquid-liquid phase separation, material properties of protein condensates, optogenetics, synthetic biology

## Abstract

Phase separation of biomolecules into condensates is a key mechanism in the spatiotemporal organization of biochemical processes in cells. However, the impact of the material properties of biomolecular condensates on important processes, such as the control of gene expression, remains largely elusive. Here, we systematically tune the material properties of optogenetically induced transcription factor condensates and probe their impact on the activation of target promoters. We demonstrate that transcription factors in rather liquid condensates correlate with increased gene expression levels, whereas stiffer transcription factor condensates correlate with the opposite effect, a reduced activation of gene expression.

We demonstrate the broad nature of these findings in mammalian cells and mice, as well as by using different synthetic and natural transcription factors. We observe these effects for both transgenic and cell-endogenous promoters. Our findings provide a novel materials-based layer in the control of gene expression, which opens novel opportunities in optogenetic engineering and synthetic biology.

## 1. Introduction

Biomolecular condensates enable the spatial and temporal organization of cellular reactions by serving as compartments that partition molecules rapidly and selectively through phase separation.^[1,2]^ Recent approaches to design biomolecular condensates have inspired a new generation of synthetic biology strategies to influence cell fate and function or to elucidate basic biological principles that rely on phase separation.^[3]^ Examples include the spatially controlled incorporation of synthetic amino acids into protein polymers,^[4]^ the control over *E.coli* biochemistry by sequestration of pathways or buffering of mRNA translation rates,^[5]^ the design of biochemical reaction centers to re-route molecular fluxes,^[6]^ the 3D organization of macromolecular complexes controlling genome architecture,^[7]^ epigenetic imprinting,^[8]^ or the modulation of gene expression.^[9,10]^

Key driving forces for biomolecular condensate formation are the concentration of the involved molecules (such as proteins, RNA, or DNA), the biophysical properties of the involved biopolymers, as well as the multivalent interactions among them. Multivalency is often given through intrinsically disordered regions (IDRs) which are stretches of amino acids without any specific secondary structure. Proteins such as FUS^[11,12]^ or HNRNPA1^[13]^ are well-known examples, described to undergo phase separation upon reaching a critical threshold concentration. Further, an increase in the multivalency of IDR interaction, for example, through post-translational modifications or mutations, can promote the transition from the liquid state to gel- or solid-like materials.^[13–18]^

Accordingly, the synthetic induction of multivalent interactions among IDRs enables the formation of biomolecular condensates on command. In a seminal study by the Brangwynne and Toettcher groups, the light-inducible homo-oligomerization of the *Arabidopsis thaliana* blue light photoreceptor Cry2 was used to increase the binding valency of IDRs and thus the formation of liquid protein condensates. By fusing IDRs to a mutant of Cry2, that forms higher-order oligomers (Cry2_olig_), light-inducible solid-like gels were obtained, as measured by fluorescence recovery after photobleaching (FRAP). Further, as a function of the protein variant and the illumination conditions, the light-responsive formation of condensates with different material properties was observed.^[19]^ Since then, light-inducible formation of condensates has been applied to elucidate fundamental mechanisms and functions of phase separation,^[19,20]^ to optimize metabolic engineering,^[6]^ or to amplify biomolecular processes such as transcription activation.^[10]^

Several recent studies have suggested that liquid-liquid phase separation (LLPS) of transcription factors plays important roles in the control of gene expression.^[21–25]^ However, the reported effects of transcription factors undergoing LLPS range from positive^[9,10,26,27]^ *via* neutral^[28]^ to negative effects^[29]^ on gene expression, with the underlying cause for this discrepancy remaining unknown. One possible mechanism that may have led to these divergent observations could be the different material properties of the investigated transcription factor condensates. Although it has been reported that material properties influence condensate function,^[30–33]^ their influence on gene expression remains inconclusive.^[34–37]^

In this study, we systematically engineered light-inducible transcription factor condensates with different material properties and analyzed their influence on transcription activation in mammalian cells and mice. We first performed an analysis of a synthetic transcription factor and subsequently applied the same concept to selected natural transcription factors of the NF-κB, STAT3, and STAT6 pathways, that play pivotal roles in important processes like the immune response. We found that transcription factor condensates that exhibit rather liquid material properties have a positive effect on transgene expression levels, whereas stiffer condensates correlated with a decrease in the expression of synthetic reporter genes or endogenous promoters. These findings provide a novel concept of how biomolecular transcription factor condensates may influence gene expression. Furthermore, the molecular tools developed here may serve as an additional, material-based layer in synthetic biology for the control of target gene regulation.

## 2. Results

### 2.1. Engineering light-inducible transcription factor condensates with graded material properties

We hypothesized that altered material properties of transcription factor condensates may impact gene expression levels. To test this hypothesis, we used our previously established optogenetically-induced transcription factor condensates. This system is based on a split synthetic transcription factor comprising the DNA-binding domain TetR fused to CIBN (CIBN-TetR) and the activation domain VP16 fused to the blue light photoreceptor Cry2, further containing eYFP for visualization (Cry2-eYFP-VP16). In this configuration, illumination with blue light triggered dimerization of CIBN and Cry2 to reconstitute a functional transcription factor.^[38]^ The incorporation of the FUS-derived IDR (FUS_N_, Cry2-eYFP-FUS_N_-VP16) led to the formation of light-inducible liquid condensates due to Cry2 homo-oligomerization. The formation of liquid condensates at the promoter site correlated with up to 5-fold increased reporter gene expression compared to the non-condensed, IDR-lacking transcription factor.^[10]^ This increase is most likely caused by the higher concentration of activation domains within the condensate in close vicinity to the promoter (Figure 1A).^[21]^

**Figure 1:**
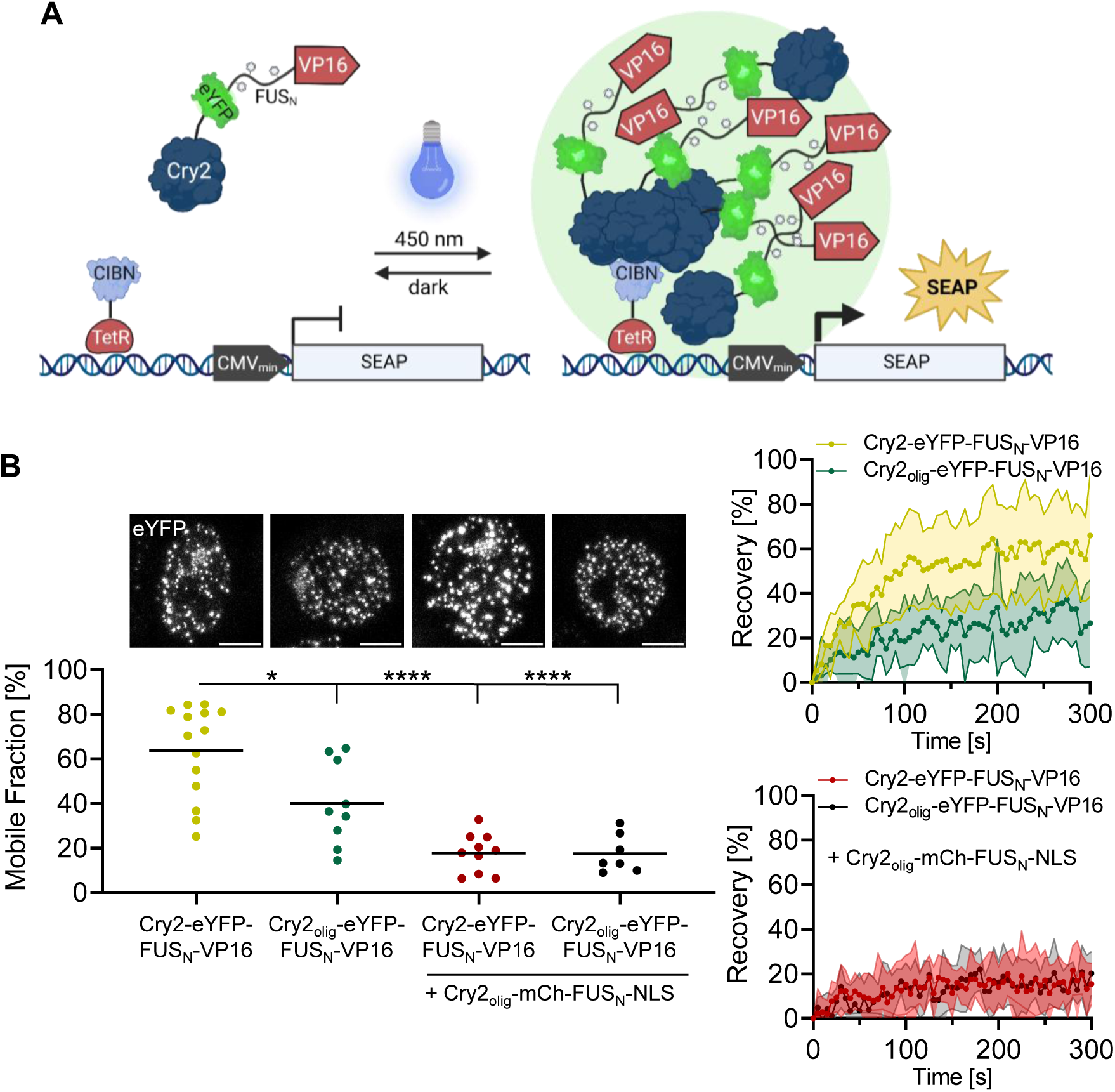
Design approach for tuning the material properties of transcription factor condensates. (**A**) Design of the light-dependent transcription factor. The transcription factor consists of two parts. First, a VP16 activation domain fused to the intrinsically disordered region (IDR) FUS_N_, eYFP for visualization, and the blue light photoreceptor cryptochrome 2 (Cry2). Second, the DNA-binding motif TetR fused to CIBN. Upon illumination with blue light, Cry2 binds CIBN and further undergoes homo-oligomerization, leading to multivalent interactions and the induction of LLPS. VP16 is recruited to the transcription start site provided by the CMV minimal promoter and induces reporter gene expression. (**B**) Tuning the material properties of transcription factor condensates. To modify condensate material properties, two strategies were pursued: first, increasing the valency of interaction by exchanging Cry2 for Cry2_olig_, which forms higher-order oligomers; and second, increasing valency and concentration by co-transfection of a construct encoding Cry2_olig_ fused to mCherry (for visualization) and FUS_N_ and an NLS. Constructs encoding CIBN-TetR and a tetO_4_-based SEAP reporter were co-transfected into HEK-293T cells together with Cry2-eYFP-FUS_N_-VP16 or Cry2_olig_-eYFP-FUS_N_-VP16 constructs (yellow and green data points). Optionally, the construct encoding Cry2_olig_-mCh-FUS_N_-NLS was added (in a 2:1 plasmid amount ratio in relation to the VP16-containing construct, red and black data points). The cells were cultivated in the dark for 32 h prior to FRAP analysis. FRAP measurements were started after 10 min of blue light illumination (2.5 µmol m^-^² s^-1^). Images show representative nuclei directly prior to droplet bleaching; scale bar = 5 µm. Graph shows the mean and single values of the mobile fractions calculated from the non-linear fits of n ≥ 7 condensate recovery curves (see right panels). Pairwise comparisons were performed using a Student’s t.test (* = *P* ≤ 0.05; **** = *P* ≤ 0.0001).

In this study, we pursued two strategies to modulate the material properties of the transcription factor condensates: first, by selecting a mutant of Cry2 that correlated with enhanced clustering (Cry2_olig_,^[39]^ Cry2_olig_-eYFP-FUS_N_-VP16) and second, by gradually increasing the concentration of IDRs by modularly adding a construct encoding additional FUS_N_ fused to Cry2_olig_ and mCherry (mCh, for visualization), by co-transfection. The construct further contained a nuclear localization sequence (NLS, Cry2_olig_-mCh-FUS_N_-NLS).

We transfected CIBN-TetR together with Cry2-eYFP-FUS_N_-VP16 or Cry2_olig_-eYFP-FUS_N_-VP16 with or without additional Cry2_olig_-mCh-FUS_N_-NLS into HEK-293T cells together with a TetR-responsive reporter construct and observed condensate formation upon blue-light illumination (Figure 1B). We then analyzed the influence of the two strategies, Cry2_olig_-enhanced clustering and increased IDR concentration, on condensate material properties by conducting FRAP experiments. We recorded the recovery of photobleached condensates (Figure 1B, Supplementary Figure 1) and compared the mobile fractions. Switching from Cry2 to Cry2_olig_ significantly decreased mobile fractions from 60% to 31%, as well as the apparent diffusion coefficient from D_app_ = 0.0026 µm^2^/s to D_app_ = 0.0017 µm^2^/s and increased the half recovery time from t_1/2_ = 42.3 s to t_1/2_ = 63.6 s. The increase in IDR concentration by the modular addition of Cry2_olig_-mCh-FUS_N_-NLS further decreased the mobile fraction to 16 or 18% for Cry2-eYFP-FUS_N_-VP16 and Cry2_olig_-eYFP-FUS_N_-VP16, respectively (Figure 1B).

Complementarily, we analyzed condensate number per cell, area, integrated optical density (eYFP), and circularity from the eYFP channel (Figure 2) and measured the mean mCherry signal in the nucleus and of the condensates to monitor expression levels of Cry2_olig_-mCh-FUS_N_-NLS as a control (Supplementary Figure 2). Both Cry2-eYFP-FUS_N_-VP16 and Cry2_olig_-eYFP-FUS_N_-VP16 yielded condensates of similar number and size. Upon addition of Cry2_olig_-mCh-FUS_N_-NLS, the condensate number per cell was significantly decreased and further diminished with increasing the amounts of this construct. Furthermore, we observed a significant increase in the condensate area, for the highest amount of Cry2_olig_-mCh-FUS_N_-NLS. The integrated optical density of eYFP (sum of all eYFP pixel values) in the condensates did not change significantly between the conditions, suggesting similar amounts of the eYFP-containing activation domains in all conditions. With increasing amounts of Cry2_olig_-mCh-FUS_N_-NLS, we observed a dose-dependent decrease in condensate circularity. Changes in condensate morphology are often associated with distinct material properties. Whereas, circularity is considered a hallmark of liquid condensate properties, gel- or solid-like condensates with larger immobile fractions can adapt more irregular, less circular shapes. The here-observed decrease in circularity is therefore in agreement with previous observations (Figure 2B).^[19,33,40,41]^

**Figure 2:**
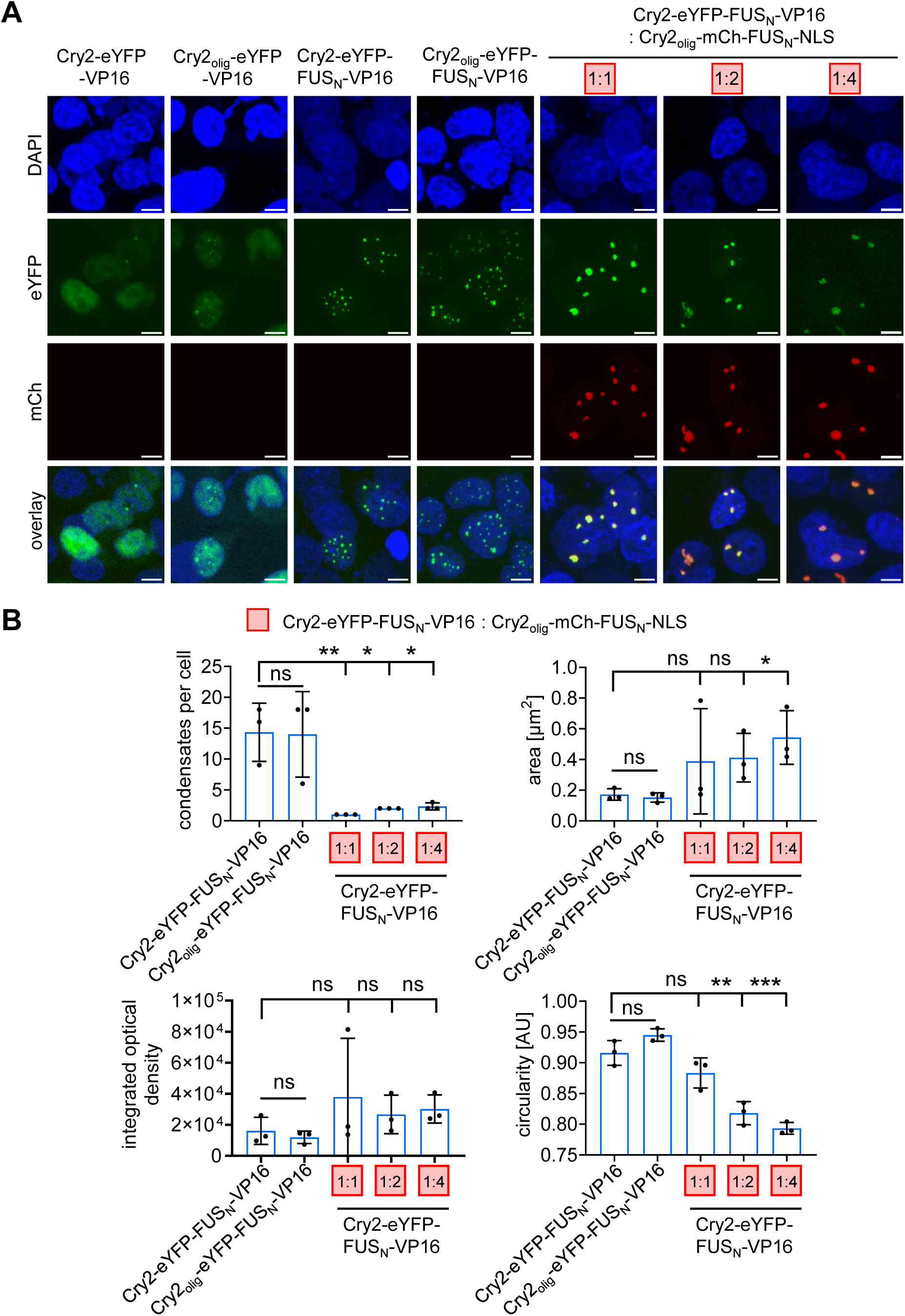
Microscopical characterization of the transcription factor condensates. (**A**) Microscopy images of the condensates. HEK-293T cells were transfected with CIBN-TetR and the indicated construct combinations together with a tetO_4_-based SEAP reporter (numbers in red boxes indicate the ratios of Cry2-eYFP-FUS_N_-VP16 to Cry2_olig_-mCh-FUS_N_-NLS constructs). 8 h post transfection, cells were illuminated with blue light for 72 h (2.5 µmol m^-^² s^-1^) and afterwards analyzed by microscopy. Representative microscopy images are shown; scale bar = 5 µm. (**B**) Quantitative analysis of the transcription factor condensates (only condensate forming conditions, panels 3-5). Number of condensates per cell, condensate area, integrated optical density (with regard to eYFP fluorescence), and circularity were determined. Single data points are the median of the population of one replicate, the bar represents the mean of n = 3 replicates. At least 215 cells or 1619 condensates were analyzed per condition. Pairwise comparisons were performed using a Student’s t.test (* = *P* ≤ 0.05; ** = *P* ≤ 0.01; *** = *P* ≤ 0.001).

### 2.2. Altered material properties of transcription factor condensates affect gene expression levels

To evaluate the effects of transcription factor condensates with different material properties on reporter gene expression, we analyzed transcription activity from a TetR-responsive promoter driving the expression of a secreted alkaline phosphatase (SEAP) as a reporter in HEK-293T cells. In this context, we first analyzed whether the condensates colocalized with the target promoter. To this end, we used a previously engineered U2-OS cell line with a chromosomal integration of an array of 256 repeats of the lacO operator in direct proximity to a tetO_96_-responsive promoter.^[42]^ We stained the lacO repeats by expressing the lacO-specific binding protein LacI fused to the blue-fluorescent protein (LacI-BFP).

In the presence of the DNA-binding domain CIBN-TetR, we observed colocalization of Cry2-eYFP-FUS_N_-VP16 and Cry2_olig_-mCh-FUS_N_-NLS with LacI-BFP, indicating that the transcription factor condensates were indeed localized at the promoter region (Figure 3A). Omission of the DNA-binding domain CIBN-TetR disrupted the colocalization of eYFP and mCh with the BFP-stained promoter region (Supplementary Figure 3A). We performed a similar experiment in HEK-293T cells using a reporter vector containing a tetO_6_ and a lacO_256_ array and confirmed the localization of Cry2-eYFP-FUS_N_-VP16 or Cry2_olig_-eYFP-FUS_N_-VP16 with Cry2_olig_-mCh-FUS_N_-NLS at the reporter plasmids (Supplementary Figure 3B).

**Figure 3:**
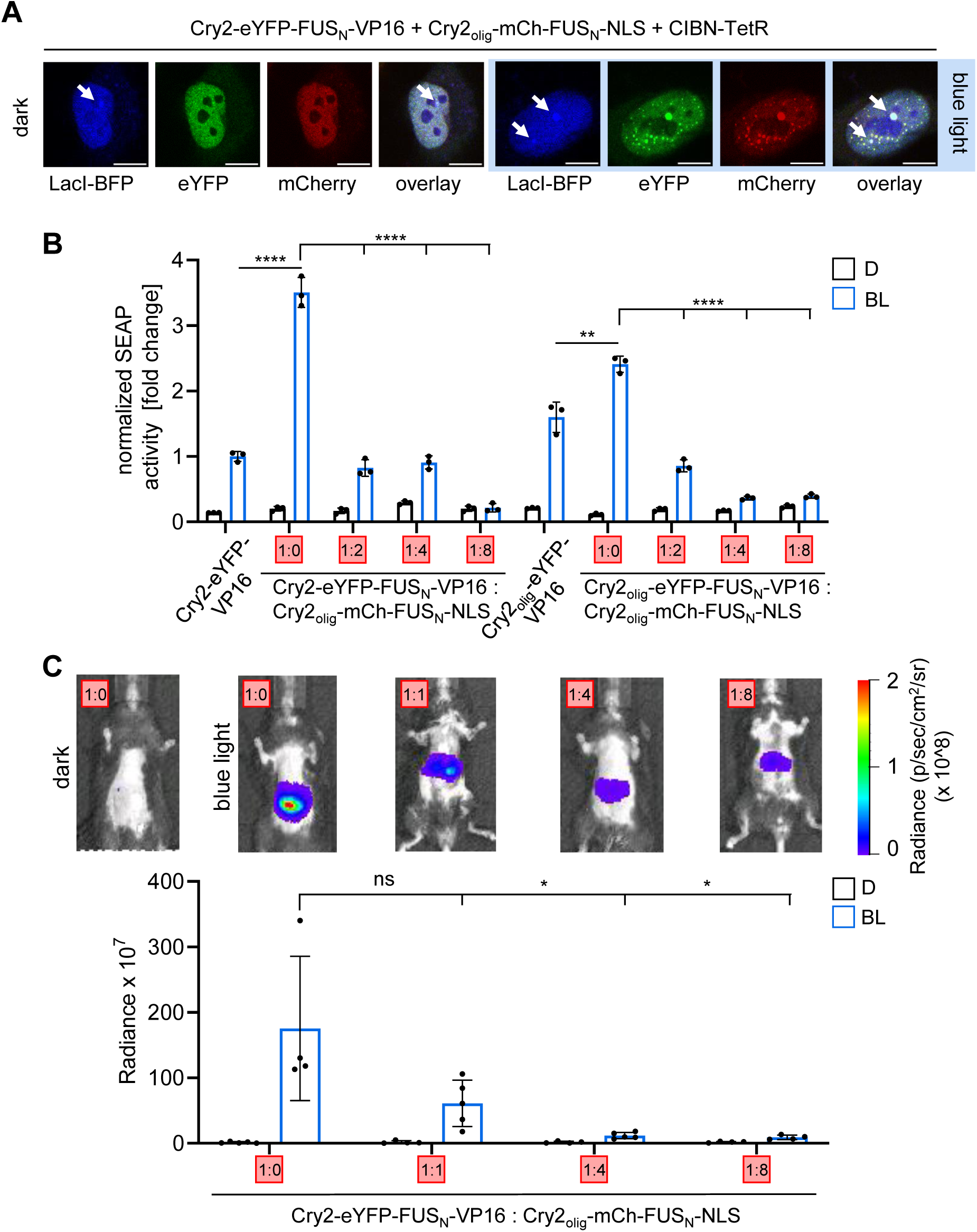
Influence of modified transcription factor condensate material properties on transcription. (**A**) Colocalization of transcription factor condensates with the target promoter. U2-OS cells harboring a genomic locus with 256 lacO and 96 tetO repeats were transfected with constructs for LacI-BFP, Cry2-eYFP-FUS_N_-VP16, Cry2_olig_-mCh-FUS_N_-NLS and CIBN-TetR. The cells were cultivated for 8 h, then illuminated for 24 h with blue light (2.5 µmol m^-^² s^-1^) and subjected to microscopy analysis. Scale bar = 10 µm. (**B**) Influence of transcription factor condensates on transgene expression. The indicated constructs were transfected into HEK-293T cells together with constructs for DNA-binding (CIBN-TetR) and a tetO_4_-based SEAP reporter. The numbers in the red boxes indicate the approximate plasmid ratio of the VP16 to mCherry-containing constructs. Cells were cultivated for 72 h in the dark (D) or under blue light (BL, 2.5 µmol m^-^² s^-1^) prior to quantifying SEAP and YFP production. SEAP activity values were normalized to the integrated eYFP fluorescence values of the dark samples. Data points represent the mean ± SD (n = 3). (**C**) Influence of transcription factor condensates on transgene expression in mice. Mice were hydrodynamically injected *via* the tail veins with constructs encoding CIBN-TetR and Cry2-eYFP-FUS_N_-VP16 (bidirectional vector), a tetO_7_-based firefly luciferase reporter, as well as the indicated ratios of Cry2_olig_-mCh-FUS_N_-NLS (red boxes, with regard to the VP16-containing construct). Eight hours after plasmid injection, the mice were exposed to blue light pulses (460 nm, 10 mW cm^-2^, 2 min on, 2 min off, alternating) for 11 h. Subsequently, luciferin was injected intraperitoneally and bioluminescence images were acquired. Mean bioluminescent radiance (p s^−1^ cm^−2^ sr^−1^) ± SEM is shown (n = 4-5) together with a representative mouse image. All pairwise comparisons were performed using a Student’s t.test (ns = *P* > 0.05; * = *P* ≤ 0.05; ** = *P* ≤ 0.01; *** = *P* ≤ 0.001; **** = *P* ≤ 0.0001).

When analyzing the effect of transcription factor condensates on gene expression, potential differences in the expression levels of the different activator constructs must be considered. To ensure that the observed effects did not derive from different transcription factor expression levels, we quantified the expression levels of our eYFP-tagged transcription factors *via* flow cytometry. The observed eYFP levels are comparable between the conditions (Supplementary Figure 3C, eYFP expression levels). We further normalized the reporter gene activity to the FACS-determined amount of transcription factor, similar to a previous study.^[10]^ In agreement with previous reports, Cry2-eYFP-FUS_N_-VP16 yielded a 3- to 4-fold increase in reporter gene expression compared to the control lacking the IDR and showing no condensate formation (Cry2-eYFP-VP16)^[10]^. Non-condensate-forming Cry2_olig_-eYFP-VP16 showed 1.6-fold higher gene expression activities compared to Cry2-eYFP-VP16, likely due to increased clustering and therefore increased VP16 recruitment. Cry2_olig_-eYFP-FUS_N_-VP16 showed a 1.5-fold increase in comparison to the corresponding non-condensate-forming, FUS_N_-devoid Cry2_olig_-eYFP-VP16 construct. Interestingly, Cry2_olig_-eYFP-FUS_N_-VP16 condensates are less active than Cry2-eYFP-FUS_N_-VP16 condensates and upon addition of increasing amounts of Cry2_olig_-mCh-FUS_N_-NLS to both conditions, blue-light-induced reporter expression was decreased step-wise, down to background levels (Figure 3B), although the amount of activation domains (as judged from eYFP fluorescence, Figure 2B) was unchanged. This suggests that stiff condensates of otherwise activating transcription factors, although bound to their promoter, correlate with reduced induction of transcription.

To analyze whether this principle is also functional *in vivo*, we transferred the system to mice. To this end, we first designed genetically more compact versions of the vector system to increase gene transfer efficiency in mice. We placed CIBN-TetR and Cry2-eYFP-FUS_N_-VP16 on a single plasmid, either separated by posttranslational cleavage sites (T2A or a combination of T2A and P2A) or under the control of two CMV promoters in either a consecutive or bidirectional configuration (Supplementary Figure 4A). We tested the constructs using a SEAP reporter under the control of a TetR-responsive promoter and chose the bidirectional configuration, as it showed the highest dark-to-blue-light fold-induction, for all subsequent experiments (Supplementary Figure 4B). We delivered the bidirectional CIBN-TetR/Cry2-eYFP-FUS_N_-VP16 construct together with a TetR-responsive luciferase reporter and increasing amounts of the Cry2_olig_-mCh-FUS_N_-NLS vector into mice *via* hydrodynamic tail vein injection. The mice were either kept in the dark or subjected to external blue light for 11 h prior to quantifying luciferase production, by whole-body bioluminescence imaging. In line with cell-culture experiments, blue light illumination activated luciferase production, which was gradually attenuated in the presence of increasing amounts of Cry2_olig_-mCh-FUS_N_-NLS (Figure 3C, Supplementary Figure 4C).

These results suggest that modulation of the material properties of synthetic transcription factor condensates at the promoter site by increasing valency and IDR concentration strongly impacts gene expression levels. While rather liquid condensates with a higher mobile fraction and faster recovery led to positive effects on reporter gene expression, stiffer, more dynamically arrested condensates (lower mobile fraction and slower recovery) correlated with attenuated reporter activation in mammalian cells and mice.

### 2.3. Impact of condensate formation on the natural transcription factor RelA

After the characterization of our synthetic transcription factor condensates, we evaluated whether the same concept of increasing or decreasing transcription activation by modulating condensate properties could be transferred to endogenous transcription factors. We chose the transcriptional activator RelA (p65), one of the core components of the canonical NF-κB signaling pathway.^[43]^

We first started with a non-optogenetic approach by fusing RelA to eYFP (eYFP-RelA) or to FUS_N_ and eYFP (FUS_N_-eYFP-RelA). We titrated the plasmid amounts used for transfection of HEK-293T cells and observed a dose-dependent increase in the fraction of cells that showed transcription factor condensates. Also, a low percentage of cells expressing high levels of eYFP-RelA showed the formation of clusters (Figure 4, Supplementary Figure 5A). We then tested the capability of the two constructs to activate transcription using an NF-κB-responsive SEAP reporter. We titrated the plasmid amounts of eYFP-RelA and FUS_N_-eYFP-RelA, quantified the eYFP expression levels *via* flow cytometry, and compared the SEAP activities of the conditions with equal eYFP and thus transcription factor expression levels. In agreement with the findings for the synthetic transcription factor, condensate formation, induced by the addition of FUS_N_, correlated with increased expression of the NF-κB-responsive SEAP reporter (Figure 4C, Supplementary Figure 5B).

**Figure 4:**
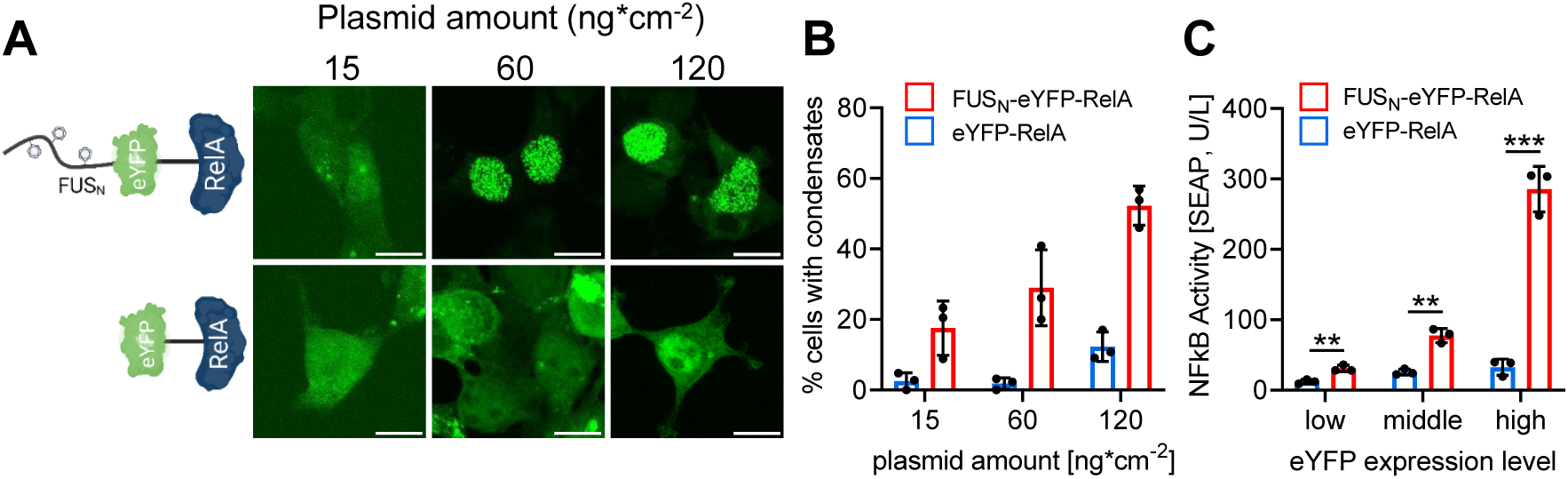
Influence of RelA-condensate formation on NF-κB-responsive reporter gene expression. (**A**) Condensate formation of RelA. Indicated amounts of expression constructs for RelA fused to eYFP and optionally FUS_N_ were transfected into HEK-293T cells together with an NF-κB- responsive firefly luciferase reporter. For internal normalization, a constitutive expression vector for renilla luciferase was co-transfected. 32 h after transfection, cells were analyzed by microscopy. Scale bar = 10 µm. Note: Minimum and maximum pixel values were adjusted differently between the panels for better visualization; quantification of the intensities is shown in Supplementary Figure 5A. (**B**) Percentage of transfected cells that form at least 1 condensate. At least 103 cells per condition were analyzed. Mean percentages of n = 3 replicates are plotted. (**C**) Influence of RelA condensate formation on reporter gene expression. HEK-293T cells were transfected with an NF-κB-responsive SEAP reporter and expression plasmids for either eYFP-RelA or FUS_N_-eYFP-RelA at increasing concentrations. 32 h after transfection, eYFP expression levels were determined via flow cytometry (see Supplementary Figure 5B). Experimental conditions with comparable eYFP expression levels (low – middle – high) were compared with regard to the amount of SEAP reporter produced. Pairwise comparisons are shown as mean ± SD (n = 3). *P* values were calculated using a Student‘s t.test: ** = *P* ≤ 0.01; *** = *P* ≤ 0.001.

### 2.4. Engineering of RelA transcription factor condensates with different material properties

Next, we evaluated the impact of light-inducible condensate stiffening on RelA activity. To this end, we fused Cry2 or Cry2_olig_ to eYFP-FUS_N_-RelA and then transfected both RelA constructs (Cry2-eYFP-FUS_N_-RelA or Cry2_olig_-eYFP-FUS_N_-RelA) together with Cry2_olig_-mCh-FUS_N_-NLS into HEK-293T cells. Cells were cultured in the dark or under blue light and subsequently analyzed by microscopy. Both constructs formed condensates in the dark, which might result from a combination of RelA’s intrinsic property to undergo LLPS^[44]^ and the IDR/Cry2-containing fusion partners (Figure 5A, B). The number of condensates per cell strongly decreased upon illumination, and an increase in condensate area was observed for the Cry2_olig_ construct (Figure 5B). These findings correspond to the findings from the synthetic transcription factor (Figure 2). The eYFP integrated optical density of the condensates (reflecting the RelA amount) was similar for all conditions; only the illuminated Cry2_olig_ constructs showed an increased value. For circularity, a decrease was observed for the Cry2_olig_ constructs similar to the synthetic transcription factor, likely reflecting stiffer condensates (Figure 5B). Next, we tested the activity of the different RelA condensates on transcriptional activation using an NF-κB-responsive dual luciferase reporter assay.^[45]^ When inducing Cry2/Cry2_olig_ clustering by blue light illumination, a strong drop in reporter gene activity was observed for both constructs. This finding is of special interest for the combination of Cry2_olig_-eYFP-FUS_N_-RelA and Cry2_olig_-mCh-FUS_N_-NLS, where the higher levels of RelA in the condensates (Figure 5B, integrated optical density) after blue-light illumination might have been expected to yield higher transcriptional activation (Figure 5C, Supplementary Figure 6A). Potential non-specific effects on reporter expression caused by the large condensates were excluded by normalizing the firefly luciferase activity with constitutively expressed renilla luciferase activity (dual luciferase assay).

**Figure 5:**
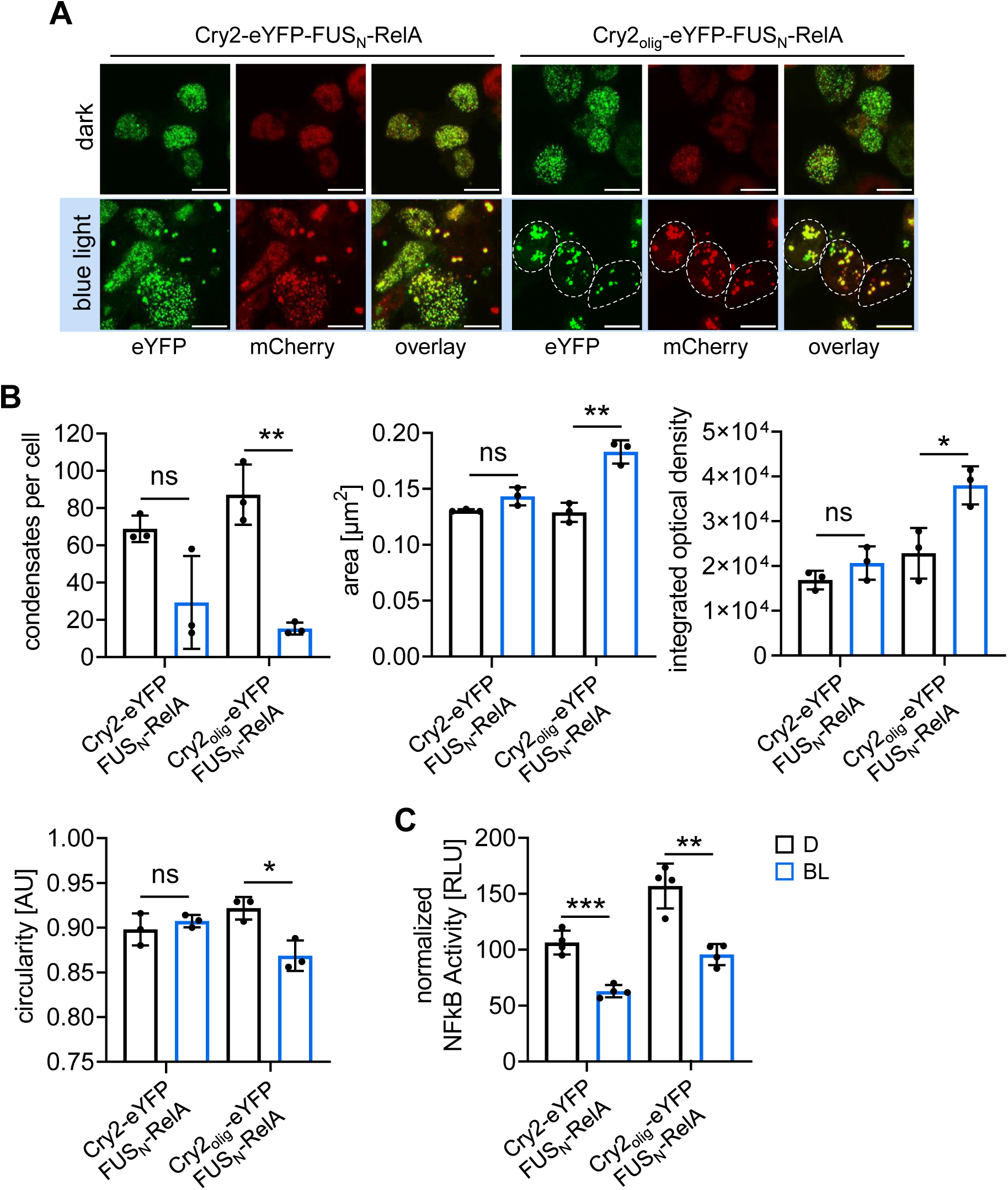
Optogenetic modulation of RelA-condensate material properties. (**A, B**) Optical characterization of RelA condensates. HEK-293T cells were transfected with Cry2/Cry2_olig_-eYFP-FUS_N_-RelA, Cry2_olig_-mCh-FUS_N_-NLS, an NF-κB-responsive firefly luciferase reporter, and a constitutive renilla luciferase reporter. 12 h post-transfection, cells were either kept in the dark or under illumination (5 µmol m^-^² s^-1^) for 40 h prior to microscopy analysis. The number of condensates per cell, condensate area, integrated optical density, and circularity were determined. At least 179 cells or 8219 condensates, were analyzed per condition of n = 3 replicates, single data points display the median of the population of each replicate. Scale bar = 10 µm. (**C**) Influence of RelA condensates on reporter gene expression. NF-κB-responsive firefly luciferase activity from the experiment conducted as described in (**A**) was quantified. Values were normalized to constitutively expressed renilla luciferase activities. Means ± SD and single values are plotted; n = 4. All pairwise comparisons were performed using a Student’s t.test (ns = *P* > 0.05; * = *P* ≤ 0.05; ** = *P* ≤ 0.01; *** = *P* ≤ 0.001).

We conclude from this data that stiffer transcription factor condensates (as induced here by the blue light-mediated multivalent interactions) correlate with reduced gene expression.

### 2.5. Downregulation of different transcription factor activities by the formation of stiff condensates

After demonstrating that stiff condensates of a synthetic or a natural transcription factor show reduced gene expression, we aimed to analyze this effect more broadly by developing a strategy that allows the recruitment of different proteins into stiff condensates. For recruitment, we chose an eGFP binding nanobody (NbGFP)^[46]^ to capture arbitrary eGFP fusion proteins. We validated this concept by fusing NbGFP to either Cry2 or Cry2_olig_ and further inserted mCherry (mCh, for visualization), FUS_N_, as well as a nuclear localization sequence (NLS, Cry2/Cry2_olig_-mCh-FUS_N_-NLS-NbGFP). We then co-transfected the construct with an eGFP-RelA expression vector and analyzed condensate formation in the dark and under blue light illumination. EGFP-RelA alone showed diffusive signals both in darkness and after blue-light illumination (Figure 6A). However, upon co-transfection with Cry2_olig_-mCh-FUS_N_-NLS and either of the NbGFP-constructs (Cry2/Cry2_olig_-mCh-FUS_N_-NLS-NbGFP), formation of large condensates was observed upon illumination. The colocalization of the mCherry and eGFP signal in the condensates further demonstrates effective recruitment of eGFP-RelA into the condensates.

**Figure 6:**
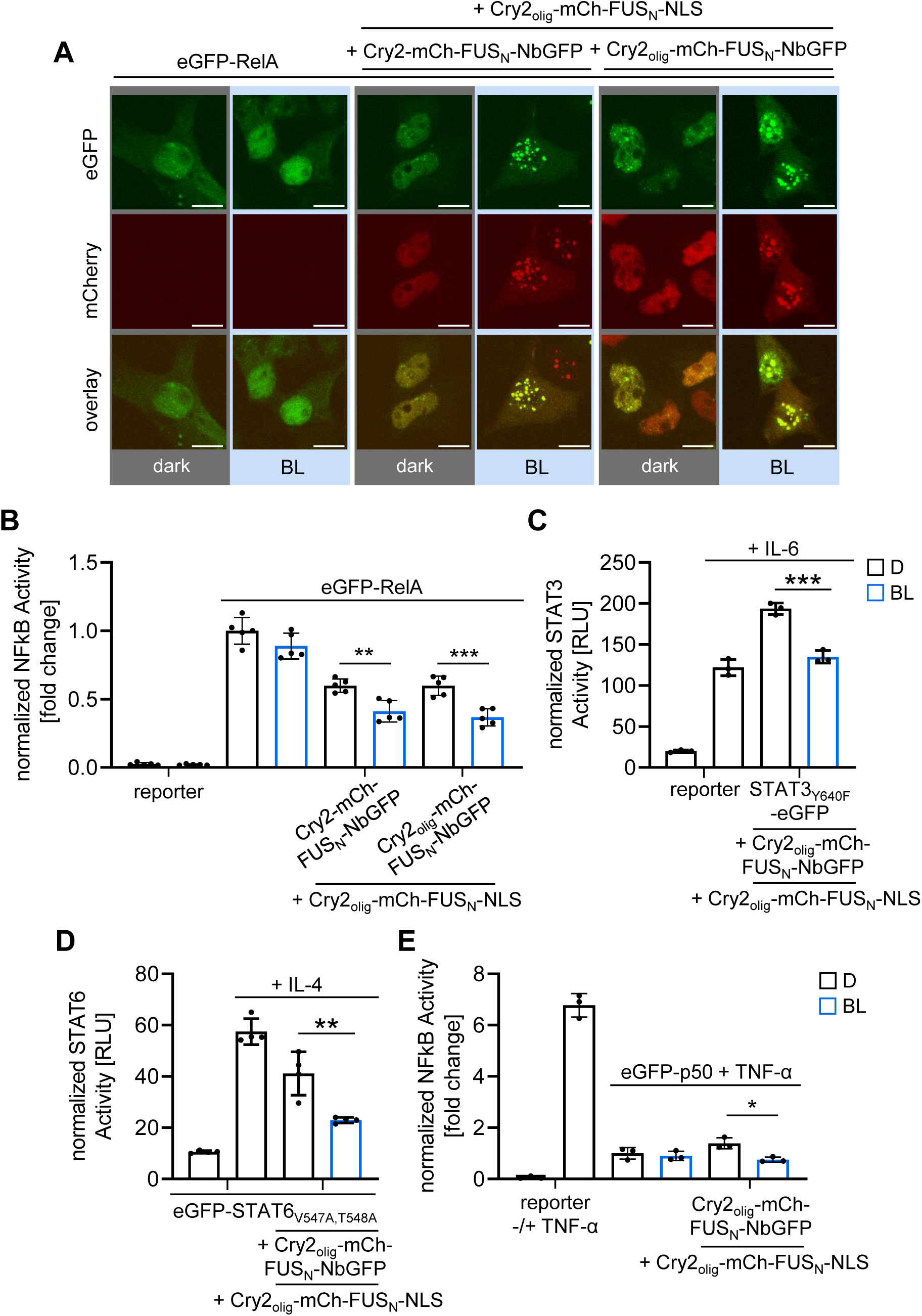
Influence of the formation of different transcription factors condensates on transcription activation. (**A**) Nanobody-mediated formation of RelA condensates. HEK-293T cells were transfected with the indicated expression vectors: an NF-κB-responsive firefly luciferase expression vector as well as a constitutive renilla luciferase construct. 8 h after transfection, cells were kept in the dark (D) or under blue light (BL) illumination (5 µmol m^-^² s^-1^) for 24 h prior to microscopy analysis. Scale bar = 10 µm. (**B**) Impact of RelA condensates on NF-κB-responsive gene expression. Cells were transfected as described in (**A**) and firefly luciferase activity was measured. Values were normalized to renilla luciferase activities. Means ± SD and single values are plotted; values are displayed as fold changes to eGFP-RelA (dark condition). (**C, D**) Impact of STAT3_Y640F_ / STAT6_V547A,T548A_ condensates on STAT3/6-responsive gene expression. The experiment was performed as described in (**A**), except that eGFP-RelA was exchanged by eGFP- STAT3_Y640F_ (**D**) or STAT6_V547A,T548A_ (**E**). Cells were stimulated with 5 ng/mL IL-6 or 10 ng/mL IL-4 for STAT3_Y640F_ and STAT6_V547A,T548A_ experiments, respectively. (**E**) Impact of p50 condensates on NF-κB-responsive gene expression. The experiment was performed as described in (**A**), except that eGFP-RelA was exchanged by eGFP-p50. The cells were further stimulated with 5 ng/mL TNF-α, 8 h after transfection. Data is displayed as fold change to eGFP-p50 (dark condition). n = 3-5, pairwise comparisons were performed using a Student’s t.test (* = *P* ≤ 0.05; ** = *P* ≤ 0.01; *** = *P* ≤ 0.001).

When analyzing luciferase expression under the control of an NF-κB-responsive promoter, a significant decrease in luciferase activity was observed under illumination compared to the dark control, with the effect being stronger for the Cry2_olig_-based construct. This finding is in line with the previous observations and demonstrates the functionality of the recruitment-based approach for a inducible decrease in RelA-responsive gene expression (Figure 6B).

Next, we aimed at validating this approach with STAT3 and STAT6 transcription factors. To challenge the system, we used mutants with increased activity (STAT3_Y640F[47]_ and STAT6_V547A/T548A[48]_) compared to the wild type. We co-transfected expression constructs for Cry2_olig_-mCh-FUS_N_-NLS-NbGFP and Cry2_olig_-mCh-FUS_N_-NLS with either STAT3_Y640F_-eGFP or eGFP-STAT6_V547A/T548A_ vectors together with a STAT3- or STAT6-responsive luciferase reporter. The cells were further stimulated with interleukin 6 (IL-6) or interleukin 4 (IL-4) to activate STAT3 and STAT6 signaling pathways, respectively. The cells were subsequently incubated in the dark or under blue light prior to measuring luciferase activity. Upon co-transfection with the condensate-forming constructs, a 1.4- and 1.8-fold decrease in reporter output was observed for STAT3_Y640F_-eGFP and eGFP-STAT6_V547A/T548A_ under blue light compared to the dark control, respectively (Figure 6C, D, Supplementary Figure 6B, C).

To confirm the binding of eGFP-RelA, STAT3_Y640F_-eGFP, and eGFP-STAT6_V547A/T548A_ to the reporter vectors, we inserted their respective response elements into the above-described reporter vector containing the lacO_256_ array. We observed colocalization of eGFP, mCherry, and BFP, suggesting that the transcription factors within the condensates are still capable of binding to DNA (Supplementary Figure 7).

Additionally, we tested the influence of stiff condensate formation on the transcriptional inhibitory p50 homodimer, which acts downstream of the canonical NF-κB signaling pathway and represses transcription when bound to its target promoter.^[49]^ We transfected HEK-293T cells with the NF-κB-responsive luciferase reporter. Stimulation with TNF-α resulted in high luciferase output. This activity was strongly reduced upon co-transfection of an eGFP-p50 construct. Upon further co-transfection with Cry2_olig_-mCh-FUS_N_-NLS-NbGFP and Cry2_olig_-mCh-FUS_N_-NLS and illumination with blue light, a further 1.8-fold decrease in reporter output was observed compared to the dark control, indicating that the formation of a stiff condensate led to further strengthened transcriptional inhibition. This further indicates the sustained binding of p50 to the promoter site after stiff condensate formation, as higher luciferase activities would have been expected if the inhibitor was detached (Figure 6E, Supplementary Figure 8).

From these observations, we conclude that optogenetically formed, stiff condensates of activating or inhibitory transcription factors can likely be used as a broadly applicable strategy to downregulate gene expression in response to light.

### 2.6. Light-controlled downregulation of endogenous promoters by stiff condensate formation

While all experiments above were performed with synthetic reporters as readout, we finally investigated whether the same effect could be observed for endogenous promoters, at the example of NF-κB-responsive promoters. To this aim, we transfected HEK-293T cells either with empty vector or eGFP-RelA or eGFP-RelA, Cry2_olig_-mCh-FUS_N_-NLS-NbGFP and Cry2_olig_-mCh-FUS_N_-NLS. The cells were cultivated under blue light or in the dark prior to extraction of total mRNA and analysis by RNAseq. For data analysis, we established a suitable threshold (| log2FC | > 0.5) and considered only genes that were deregulated more strongly. This eliminated genes that were only marginally deregulated, for example, due to minor differences in the cultivation conditions in darkness and blue light (Table S1, Supplementary Figure 9). Next, we filtered for genes that were previously reported to be induced by NF-κB signaling^[50]^ and therefore could be bound and activated by eGFP-RelA, and affected by the condensate formation at their promoter. This step should exclude genes that were not bound by eGFP-RelA but are secondarily deregulated, for example, due to deregulation of eGFP-RelA targets that are positive or negative regulators themselves (Table S1). With this method, we obtained 15 genes that were significantly deregulated between the empty vector and eGFP-RelA conditions AND deregulated between darkness and blue light of the condensate condition (eGFP-RelA, Cry2_olig_-mCh-FUS_N_-NLS-NbGFP and Cry2_olig_-mCh-FUS_N_-NLS, Supplementary Figure 10, Supplementary Figure 11).

Indeed, all 15 genes were upregulated by eGFP-RelA and downregulated by condensate formation (Figure 7). Furthermore, we confirmed the deregulation of three of these genes (*TNFAIP3, NFKBIA, and CXCL1*) via qPCR and observed an up to 4.4-fold decrease in endogenous gene expression after condensate formation under illumination compared to the dark control (Supplementary Figure 12). The number of 15 specifically deregulated NF-κB target genes was lower than expected, which might result, for example, from cell line specifics or compensatory mechanisms of endogenous feedback mechanisms^[51,52]^.

**Figure 7:**
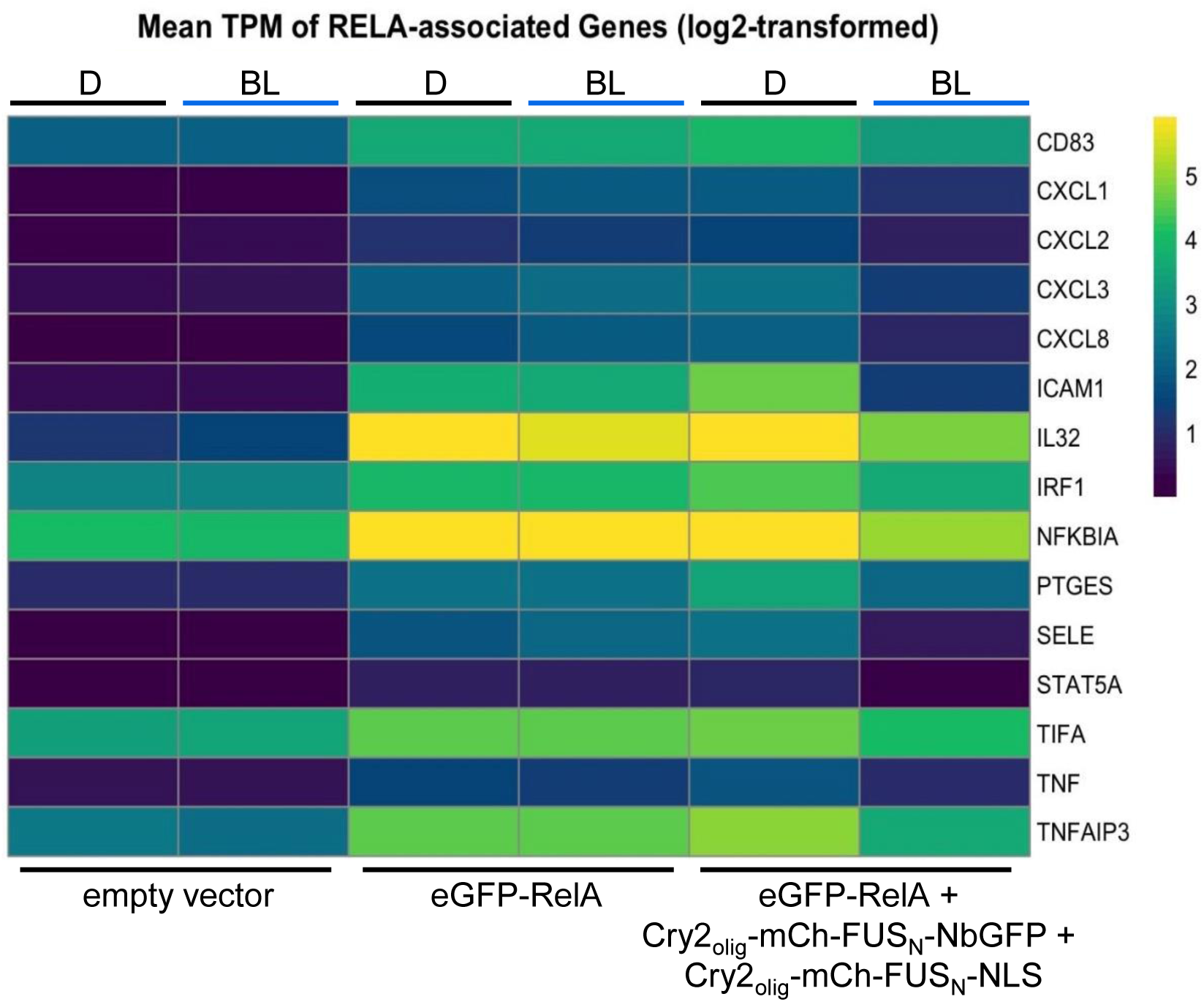
Impact of stiff eGFP-RelA condensates on endogenous gene expression. HEK-293T cells were transfected with the indicated expression vectors and an NF-κB-responsive SEAP reporter. 8 h after transfection, cells were either kept in the dark (D) or under blue light illumination (BL, 5 μmol m^-^² s^-1^) for 24 h prior to RNA extraction. Total RNA samples were subjected to RNAseq. Datasets of triplicates were pooled and 15 direct targets of eGFP-RelA were identified. Genes with a q-value ≤ 0.05 and | log2FC | > 0.5 were considered significantly deregulated. The heatmap shows their log2-transformed, mean transcripts expression levels in units of kilobase million (TPM).

In conclusion, these findings further corroborate the above-described data and support the hypothesis that optogenetically induced stiff transcription factor condensates at promoter sites lead to light-inducible downregulation of transcription factor activity at synthetic and endogenous promoters (Figure 7).

## 3. Conclusion

Recent gene activation models suggest important roles for phase separation in the control of gene expression.^[21,53]^ The IDRs of cofactors, transcription factors or polymerase II mediate phase separation and thereby, effective recruitment and concentration of cofactors and activators into transcriptional condensates.^[54–56]^ Furthermore, condensates are capable of reconfiguring chromatin interactions to access enhancer regions through DNA looping, enabling longrange communication.^[57,58]^ However, the functional impact of transcriptional condensates on gene expression levels remained inconclusive. Some groups reported positive effects^[9,10,26]^ likely due to a higher concentration of transactivation domains at promoter regions^[27]^ or increased recruitment of activating cofactors.^[56]^ Other groups reported no additional effect in comparison to a multivalent but non-phase-separating transcription factor^[28]^ or an optimum of IDRs at the promoter to achieve maximum expression levels, whereas higher IDR levels that led to condensate formation had negative effects.^[29,59]^ Further studies suggested mechanisms such as the reduced capability of transcription factor IDRs to recruit mediators due to changes in their sequence,^[60,61]^ recruitment of proteins that alter the surface tension and therefore reduce DNA binding capacity^[62,63]^ or sequestration of transcription factors in the cytoplasm^[64]^ to be involved in the downregulation of gene expression.

To investigate whether changing the material properties of transcription factor condensates could impact gene expression levels, we developed an optogenetic strategy to induce transcription factor condensates and modified their material properties by the modular addition of Cry2_olig_-mCh-FUS_N_-NLS, which increased the IDR concentration and might act as a multivalent scaffold in line with a very recent study by Hernandez-Candia *et. al.*^[33]^ We characterized their material properties and morphology and linked them to different levels of gene expression. We observed that liquid condensates have a positive effect on gene expression in comparison to diffusive transcription factors, whereas stiffer promoter-bound transcription factor condensates result in reduced transcription. We linked functional properties to certain morphological hallmarks: stiffer condensates correlated with lower circularity, a larger area, and a lower number per cell. Inhibition of gene expression occurred despite the equal or increased transcription factor amounts within the condensates, as indicated by the fluorescent intensities of the condensates. The data on the synthetic TetR-VP16-based droplet-transcription factor and the RelA-condensates suggested that rather liquid condensates led to increased gene expression levels in comparison to diffuse transcription factors, whereas stiff, more dynamically arrested condensates reversed this effect and led to reduced transcription activation.

The colocalization study (Figure 3A, Supplementary Figure 3A, B, Supplementary Figure 7) confirmed the localization of the condensates at the promoter. The smaller changes in the system, like the mutation of Cry2 to Cry2_olig_, resulted in reduced gene expression but did not lead to morphological changes (Figure 3B, difference between Cry2-eYFP-FUS_N_-VP16 and Cry2_olig_-eYFP-FUS_N_-VP16-induced reporter gene expression, although no changes in condensate morphology or condensate number were observed (Figure 2)). A possible reason for the reduced transcriptional activity of Cry2_olig_-eYFP-FUS_N_-VP16 could be the decreased dynamics and apparent diffusion coefficient that were measured by FRAP (Figure 1).^[30,31]^ The larger morphological changes that occurred with Cry2_olig_-mCh-FUS_N_-NLS addition might facilitate additional inhibitory mechanisms, such as blocking the access of activating factors to endogenous promoters, due to their larger size.

Light is a powerful stimulus to modify protein interactions non-invasively,^[65–67]^ and the incorporation of the eGFP-binding nanobody enabled us to more broadly apply our system by facilitating the incorporation of arbitrary eGFP-tagged proteins into biomolecular condensates. The constructs can be used to probe protein behavior in different phase-separated conditions by modular modification of the condensate material properties, such as exchanging Cry2 with Cry2_olig_ or adding Cry2_olig_-mCh-FUS_N_-NLS.

In conclusion, we present a designer approach for the modulation of the material properties of biomolecular transcription factor condensates. We link them to distinct biological outputs using the example of transcriptional activation. We believe that the rational design of biomolecular transcription factor condensates with defined properties will enable new avenues that allow for controlling and tuning of cellular processes or the implementation of new functions. The approach presented here is likely transferable to probe and control the function of other proteins and processes in different directions of fundamental and applied research.

## 4. Experimental Section

### 4.1. Cloning

All plasmids in this study were cloned *via* Gibson Assembly^[68]^ or Aqua Cloning^[69]^ and are described in Table S1. Mutations and short sequences, such as linkers, were inserted with oligonucleotides and PCR. For constructs that contain multiple repeats (promoter response elements and double vectors containing two CMV promoters), *E.coli* were grown at room temperature or a maximum of 30 °C. All NF-κB and STAT3-related reporters and templates for PCR amplification of NF-κB and STAT3 signaling proteins were a kind gift from the laboratory of Bodo Grimbacher (Institute for Immunodeficiency, Freiburg). All sequences were confirmed by Sanger sequencing.

### 4.2. Illumination

All optogenetic experiments in 96-well format were conducted in black clear-bottom plates (µCLEAR®, Greiner Bio-One, catalog no. 655090) and illuminated using the optoPlate-96^[70]^ equipped with 470 nm LEDs (Würth Elektronik, MPN: 150141RB73100) that was programmed using the optoConfig-96^[71]^. Experiments in 24- or 6-well format were illuminated with micro- controller-regulated illumination panels containing LEDs with 460 nm (LED Engin, MPN: LZ1-10B202-0000). For live cell imaging, illumination was conducted directly at the confocal microscope using a pE-4000 LED light source (460 nm, CoolLED, Andover, UK). All samples that express photosensitive proteins were handled under dim, green, safe light to avoid unwanted photoreceptor activation.

### 4.3. Cell culture and transfection

HEK-293T (DSMZ, catalog no. ACC 635) and U2OS 2-6-3^[42]^ cells were cultivated in Dulbecco’s modified Eagle’s medium (DMEM, PAN Biotech, catalog no. P04-03550) completed with fetal calf serum (FCS, 10% (v/v), PAN Biotech, catalog no. P30-3602) and penicillin-streptomycin (1% (v/v), PAN Biotech, catalog no. P06-07100) at 37 °C and 5% CO_2_. They were passaged every 2-3 days after reaching a confluency of ≈ 80%. For experiments, cells were seeded into multi-well plates and transfected with polyethyleneimine (PEI, 1 mg/mL in H_2_O, pH 7, Polyscience, catalog no. 23966-1). Therefore, DNA and PEI were prepared separately, each in half of the total volume Opti-MEM (Thermo Fisher Scientific, catalog no. 22600-134). The two approaches were mixed, immediately vortexed for 15 s, and incubated for 15 min at room temperature. DNA/PEI mixes were then applied dropwise to the cells. Cells transfected for optogenetic experiments were then kept in darkness. If indicated, cells were stimulated 8 h after transfection with TNF-α (Merck, catalog no. H8916-10UG), IL-6 (Prepotech, catalog no. 200-06) or IL-4 (produced as previously described^[72]^). Concentrations are indicated in the figure legends. Exact plasmid combinations and amounts for each experiment are listed in Table S2.

### 4.4. Reporter assays

All reporter assays (SEAP and dual-luciferase) were performed in 96-well plates (non-optogenetic experiments: Corning, catalog no. 3596; optogenetic experiments: µCLEAR®, Greiner Bio-One, catalog no. 655090). 15,000 cells per well in 100 µL DMEM were seeded 20-24 h prior to transfection. For transfection, 125 ng DNA and 0.41 µL polyethyleneimine in 20 µL Opti-MEM per well were used. For optogenetic experiments, illumination was started 8-12 h after transfection and the assays were conducted 24-72 h after transfection, as indicated for each experiment. For the readout, either a Synergy 4 multimode microplate reader (BioTek Instruments Inc.), an Infinite 200Pro microplate reader (Tecan Trading AG), or a SpectraMax iD5 microplate reader (Molecular Devices GmbH) was used.

For firefly/renilla dual luciferase assays, cells were lysed by addition of 100 µL lysis buffer (25 mM Tris/HCl, pH 7.8, 1% Triton X-100 (v/v), 15 mM MgSO_4_, 4 mM ethylene glycol tetraacetic acid (EGTA), 1 mM DTT) per well. After 5 min of incubation at room temperature (RT), cells were resuspended and split into 2x 40 µL samples by transferring them into two white flat-bottom 96-well plates (Corning Incorporated, catalog no. CORN3912). The plates were spun down for 30 s at 1200 rpm to remove all bubbles. One plate was used for Firefly luciferase measurement by addition of 20 µL firefly substrate (0.47 mM luciferin in 20 mM Tricine, 2.67 mM MgSO_4_, 0.1 mM EDTA, 33.3 mM DTT, 0.52 mM ATP, 0.27 mM Acetyl-CoA, 5 mM NaOH, 0.264 mM MgCO_3_) per well and the other plate was used for renilla luciferase measurement by addition of 20 µL coelenterazine (stock: 472 µM coelenterazine in methanol; diluted 1:10 in PBS directly before measurement) per well. Both plates were measured immediately after substrate addition (program: 10 s shaking, 1000 ms integration time, endpoint measurement). Firefly values were normalized with background-subtracted renilla values.

For SEAP activity measurement, the whole supernatant was transferred into a U-bottom transparent 96-well plate (Carl Roth, catalog no. 9291.1), sealed with adhesive tape, and heated at 65 °C for 30 min. Afterwards, samples were spun down for 3 min at 1250 rpm, then 80 µL supernatant was added to 100 µL 2x SEAP buffer (21% (v/v) diethanolamine, 20 mM L-homoarginine-hydrochloride, 1 mM MgCl_2_ (pH 9.8)) in transparent flat-bottom 96-well plates (Carl Roth, catalog no. 9293.1). Directly prior to the measurement, 20 µL para-nitrophenyl phosphate solution (pNPP, 120 mM) was added and its conversion to para-nitrophenol by SEAP was measured by absorbance at 405 nm in time intervals of 1 min. SEAP activity was calculated as described previously.^[73]^

### 4.5. Flow cytometry

For comparison between different constructs in reporter assays, the expression levels of the fluorescent proteins in each construct were determined by flow cytometry. Therefore, medium was removed and cells were detached with 50 µL trypsin/EDTA solution (PAN Biotech, catalog no. P10-023500) per well. Afterwards, 150 µL PBS supplemented with FCS (2% (v/v)) was added and cells were carefully resuspended and transferred into a transparent U-bottom 96-well plate (Greiner, catalog no. 650161). Cells were kept on ice until measurement with an Attune NxT flow (Thermo Fisher Scientific). EGFP and EYFP were measured by excitation with a 488 nm laser and detected with a 530/30 nm emission filter (BL1) and mCherry was measured with a 561 nm excitation laser and a 620/15 nm emission filter (YL2). 15,000 cells per well were analyzed at a flow rate of 200 µL/min. Analysis was performed using the FlowJo_v10™ software. To obtain the expression levels of the eYFP-tagged transcription factor, singlets were gated and then the integrated eYFP intensity (count of singlet population multiplied by the mean eYFP fluorescence intensity of the singlet population) was calculated and used for normalization.^[10]^

### 4.6. Microscopy and image analysis

For microscopy of fixed samples, coverslips (Carl Roth, catalog no. YX03.2) were placed in 24-well plates (Corning, catalog no. CORN3524) and coated with 500 µL rat tail collagen I (Thermo Fisher Scientific, catalog no. A1048301) at a concentration of 50 µg/mL diluted in 25 mM acetic acid. After one hour of incubation at RT, the wells were washed three times with 500 µL PBS, and 75,000 cells per well were seeded. For transfection in 24-well format, 750 ng DNA and 2.4 µL PEI in 100 µL OptiMEM per well were used. Illumination of optogenetic experiments was started after 8 h for time periods as indicated in the figure legends. Cells were fixed by removing the medium and adding 200 µL methanol-free paraformaldehyde (PFA, Science Services, catalog no. E15714-S; 4% diluted in PBS (v/v)) under green safe light. After 15 min of incubation at RT in darkness, PFA was removed and cells were washed twice with 500 µL PBS. For staining of the nuclei, 500 µL 0.2 µg/mL 4’,6’-diamidino-2-phenylindole (DAPI, Merck, catalog no. D9542) in PBS was added for 15 min at RT, then cells were washed again twice with 500 µL PBS and mounted on microscopy slides with 8.5 µL Mowiol mounting medium (2.4 g Mowiol, 6 g glycerol, 6 mL of H_2_O and 12 mL of Tris/HCl (pH 8.5)). After the samples were dried, the edges were additionally fixed with transparent nail polish. All images and time series were acquired with a Zeiss LSM 880 laser scanning confocal microscope using a 63x Plan-Apochromat oil objective (NA 1.4) in z-stacks with a distance of 1 µm. DAPI was imaged with the 405 nm laser, eGFP and eYFP were excited with the 488 nm laser, and mCherry with the 561 nm laser. For colocalization experiments, only one focal plane is shown.

For quantitative image analysis, at least 2 images of 3 replicates were acquired. Z-stacks were reduced to two-dimensional maximum intensity projections. The percentage of cells that form droplets was determined by first segmenting and counting the transfected cells and then manually counting the cells with droplets. For analysis of the droplet properties, nuclei and droplets were segmented and analyzed using custom-written macros for ImageJ 2.3.0/1.53q including the Biovoxxel Toolbox v2.6.0.^[74]^ A threshold for minimum 1 droplet per cell with a size of at least 0.017 µm^2^ (7 pixel) was set. For droplet intensity, the RawIndDen (sum of all pixel values of the droplet) was used. All macro outputs were manually double-checked. If not stated otherwise, minimum and maximum pixel values of a channel were adjusted equally for all samples of an experiment. In overlay images, intensities may be adjusted differently for better visualization of all channels.

FRAP measurements were conducted in 35 mm glass bottom dishes (µ-Dish 35 mm, high, Ibidi cat. No. 81156) using a Tokai Hit stage top incubator under continuous illumination with the pE-4000 CoolLED (460 nm, 5 µmol m^−2^ s^−1^). Time series of at least 300 s were acquired as z-stacks of nine focal planes with a distance of 600 nm, an interval of every 5 s, and the zoom factor six. Four eYFP droplets were selected simultaneously with regions of interest (ROI) of ≈ 0.8 µm diameter and bleached after the first z-stack was acquired, using 100% laser power of the 488 nm laser. For analysis, droplets were manually tracked with a circular mask of constant diameter in the x, y, and z directions for 300 seconds. Afterwards, background signals were subtracted and droplet recovery was corrected for photobleaching by normalization with the linear regression of the average fluorescence signal decrease of the nuclei. Values were normalized to the intensity before and directly after bleaching. Mobile fractions were calculated by fitting the recovery curves to an exponential plateau equation (**Y=Y_M_ -(Y_M_-Y_0_)*exp^(-k*x)^**, Y_M_ = maximum population/mobile fraction, Y_0_ = starting population, k = time rate constant) using GraphPad Prism 9.2.0. Half-maximum recovery time (t_1/2_) and the apparent diffusion coefficient were calculated from the curve fits of the mean recovery curves, with the formulas **t_(1/2)_ = ln(2)/k** and **D_app_=r²_bleach_/(1/k)** with a bleaching radius **r_bleach_** = 0.4 µm, as previously reported.^[35,54]^ Please note, that we did not calculate **D_app_** for curves that showed less than 20 % recovery and therefore reflect extremely low dynamics leading to unreliable k values in the curve fit.

### 4.7. RNA extraction and reverse transcription quantitative PCR (RT-qPCR)

For RNA extractions, 600,000 cells/well were seeded onto 6-well plates (Corning, catalog no. CORN3516) and 24 h later transfected with 600 µL transfection mix containing 3.750 µg total DNA and 12.375 µL PEI per well. Blue-light illumination (460 nm, 5 µmol m^-2^ s^-1^) was started 8 h after transfection. Cells were harvested 24 h later using 1 mL TriFast™ (VWR, cat. no. 30-2010) per well. Total RNA was extracted according to the manufacturer’s protocol. RNA integrity was verified on an agarose gel and concentrations, 260/280 and 230/280 ratios were determined with a nanodrop 1000 (Thermo Fisher Scientific).

Reverse transcription was performed using the High Capacity Kit (Applied Biosystems™, cat. no. 4368814) according to the manufacturer’s protocol using 2 µg RNA. The resulting cDNA was diluted 1:20 and used for qPCR. The primers for the three NF-κB target genes were selected from literature (NFKBIA, oAF469: 5’-ATGTCAATGCTCAGGAGCCC-3’ and oAF470: 5’-GACATCAGCCCCACACTTCA-3’;^[75]^ TNFAIP3, oAF467: 5’- CGTCCAGGTTCCAGAACACCATTC-3’ and oAF468: 5’- TGCGCTGGCTCGATCTCAG- TTG-3’;^[76]^ CXCL1, oAF471: 5’-AACCGAAGTCATAGCCACAC-3’ and oAF472: 5’-GTT-

GGATTTGTCACTGTTCAGC-3’^[77]^) and GUS was used as housekeeping gene (oMH703/oAF481, 5’-CGTCCCACCTAGAATCTGCT-3’ and oMH702/oAF482, 5’-TTGCTCACAAAGGTCACAGG-3’). qPCRs were conducted with a CFX384 Touch Real-Time PCR Detection System (BioRad) in 10 µL approaches with the PowerTrack™ SYBR Green Mastermix (Applied Biosystems™, cat. no. A46110) according to the manufacturer’s protocol (fast cycling mode with default dissociation step). ΔΔCt analysis was performed with the means of three technical replicates and the housekeeping gene GUS. Data is displayed as fold-changes to the eGFP-RelA containing control. Single data points display the biological replicates.

### 4.8. RNAseq

#### 4.8.1 Sample Preparation

Experimental procedure and total RNA extraction was performed as described in section 4.7. Additionally, RNA was cleaned up using an RNeasy Mini Kit (Qiagen, cat. no. 74104) according to the manufacturer’s protocol, including an on-column DNase I digest using 50 µL reactions of DNase I (New England Biolabs, cat. no. M0303). RNA integrity was verified at BGI Tech Solutions (Hong Kong) Co. using a Bioanalyzer (Agilent 2100 Bioanalyzer).

#### 4.8.2 Pre-Processing

RNA-seq libraries from the 18 samples were sequenced by BGI Tech Solutions on a DNBSEQ platform using paired-end chemistry with a read length of 100 base pairs each. Each strand was sequenced across two separate lanes, generating a total of 4 FASTQ files per sample. FASTQ files were obtained from BGI and subsequently quality-controlled using FASTQC^[78,79]^.

After verifying that there were no quality issues specific to individual lanes, the reads from the strand-specific sequencing replicates were pooled together. This was done to improve the accuracy and depth of coverage. After this, samples had two FASTQ files each – one for each strand.

FASTQC reported < 0.1% adapter contamination across all samples. Cutadapt^[80]^ was used for adapter trimming. Quality trimming was done using Q=30 as a threshold to filter out low-quality bases using bbduk.sh from the BBMap suite^[81]^. Length filtering was done using reformat.sh (BBMap) to filter out reads shorter than 50 bp.

#### 4.8.3 Quantification and DGE Analysis

Transcript quantification was done using Kallisto^[82]^. The reference transcriptome was obtained from Gencode (Release 38)^[83]^. Sleuth ^[84]^ was used to process the transcript abundances generated by Kallisto to perform differential gene expression analysis. By setting the gene_mode parameter to TRUE while creating a sleuth object, the quantification data from the transcript level was aggregated to the gene level to perform gene-level differential expression analysis. Sleuth offers two methods to test for differential expression. The default, Likelihood Ratio Test (LRT), assesses the probability of the data under the “full” and “reduced” models. The full model assumes that gene abundances are influenced by exposure to blue light or darkness. The reduced model assumes that the treatment has no effect on gene abundance. After estimating the ratio of the two likelihoods, it returns a test statistic with its p-value. Genes with FDR-adjusted p-value < 0.05 were considered significantly differentially expressed. The Wald Test uses the full model to test whether the coefficients associated with the conditions are significantly different from zero. In this context, each coefficient represented the effect of a specific condition (darkness/ blue light exposure) on the expression of the gene. The coefficients can be treated as estimates of fold changes. To enhance the robustness of the analysis, we designated the genes detected as significantly differentially expressed by both the LRT and Wald tests as true positives. For instance, in the volcano plots, the true positive deregulated NF-κB target genes^[50]^ are marked in red. The orange points (genes) above the significance threshold were considered false positives since they were reported by the Wald Test as significantly differentially expressed but not by the LRT.

#### 4.8.4 Identification of direct targets of eGFP-RelA

A large list of deregulated genes was identified (Table S1). To distinguish between direct and indirect NF-kB targets, this list was compared to a list of 409 NF-kB target genes compiled by the Gilmore lab.^[50]^ This list includes target genes of the NF-kB family and genes that have a kB domain in the promoter region for potential NF-kB binding but have not directly been shown to be targeted. Furthermore, genes with a low deregulation log2FC (| log2FC | < 0.5) were excluded to filter out potential effects that derive from illumination or that are subject to compensatory effects.

### 4.9. Mouse experiments

All the animals were obtained from the ECNU Laboratory Animal Centre, kept on a standard alternating 12-h light/12-h dark cycle, and were given a normal chow diet [6% fat and 18% protein (wt/wt)] and water. Wild-type C57BL/6 mice (6-week-old, male) were randomly divided into four groups. The mice were hydrodynamically injected in their tail vein with a total amount of 330 µg in 2 mL (10% of the body weight in grams) of Ringer’s solution (147 mM NaCl, 4 mM KCl, and 1.13 mM CaCl_2_) within 3-5 s. Plasmid combinations are described in Table S2. Eight hours after injection, the mice were exposed to blue light pulses (460 nm, 10 mW cm^-2^, 2 min on, 2 min off alternating; Shenzhen Kiwi Lighting Co. Ltd.) for 11 h. The control mice were kept in the dark. For in vivo bioluminescence imaging, each mouse was intraperitoneally injected with luciferin substrate solution (150 mg/kg; luc001, Shanghai Sciencelight Biology Science & Technology) under ether anesthesia. Five minutes after luciferin injection, bioluminescence images of the mice were acquired using the IVIS Lumina II in vivo imaging system (Perkin Elmer, USA). Radiance (p s^-1^ cm^-2^ sr^-1^) values were calculated for the region of interest using the Living Image® 4.3.1 software.

### 4.10. Software

All analyses were performed with Microsoft Excel 2016 and GraphPad Prism 9.2.0. FACS data was analyzed with FlowJo_v10™ and image analysis was done with ImageJ 2.3.0/1.53q including the Biovoxxel Toolbox v2.6.0^[74]^. The bioluminescent mouse images were analyzed with the Living Image® 4.3.1 software. VENN diagrams were generated using the InteractiVenn tool^[85]^. All schemes were created with BioRender.com.

## Ethics

The experiments involving animals were approved by the East China Normal University (ECNU) Animal Care and Use Committee and in direct accordance with the Ministry of Science and Technology of the People’s Republic of China on Animal Care guidelines. The protocol (protocol ID: m20221109) was approved by the ECNU Animal Care and Use Committee. All animals were euthanized after the termination of the experiments.

## Author contributions

A.A.M.F., D.K., M.M.G., J.G., and J.G. have performed experimental work. A.A.M.F., H.B.R., C.B., and W.W. analyzed the results. A.A.M.F. and W.W. wrote the manuscript. M.F. and B.G. contributed to the design of the study and in data interpretation. H.Y., V.H., F.L., and W.W. supervised the work. All authors edited and approved the manuscript.

## Data and materials availability

All data needed to evaluate the conclusions in the paper is present in the paper and/or Supplementary Materials. Raw data (images, FACS data, SEAP, and Luciferase measurements), plasmids, and detailed plasmid maps are available from the authors upon request. RNAseq data and metadata will be uploaded to the European Nucleotide Archive.

## Acknowledgmen**ts**

We are very grateful to M. Klenzendorf and D. Gaspar for their excellent technical assistance and to L. Qiao for excellent assistance in the animal work. We thank all members of the Weber laboratory for their helpful comments, especially N. Schneider, M. Hörner, and O. Thomas for valuable advice. We would like to thank the Faculty of Biology technical workshop for the construction of the illumination devices. We acknowledge the excellent scientific and technical assistance of the Signalling Factory Core Facility staff of the Albert-Ludwigs-University Freiburg for help on flow cytometry, especially P. Salavei and N. Gensch. We acknowledge the staff of the Life Imaging Center (LIC) in the Center for Biological Systems Analysis (ZBSA) of the Albert-Ludwigs-University Freiburg for their help with microscopy resources and their excellent support in microscopy setup and image acquisition.

Received: ((will be filled in by the editorial staff))

Revised: ((will be filled in by the editorial staff))

Published online: ((will be filled in by the editorial staff))

