## Supplementary Materials for "Engineering Material Properties of Transcription Factor Condensates to Control Gene Expression in Mammalian Cells and Mice"

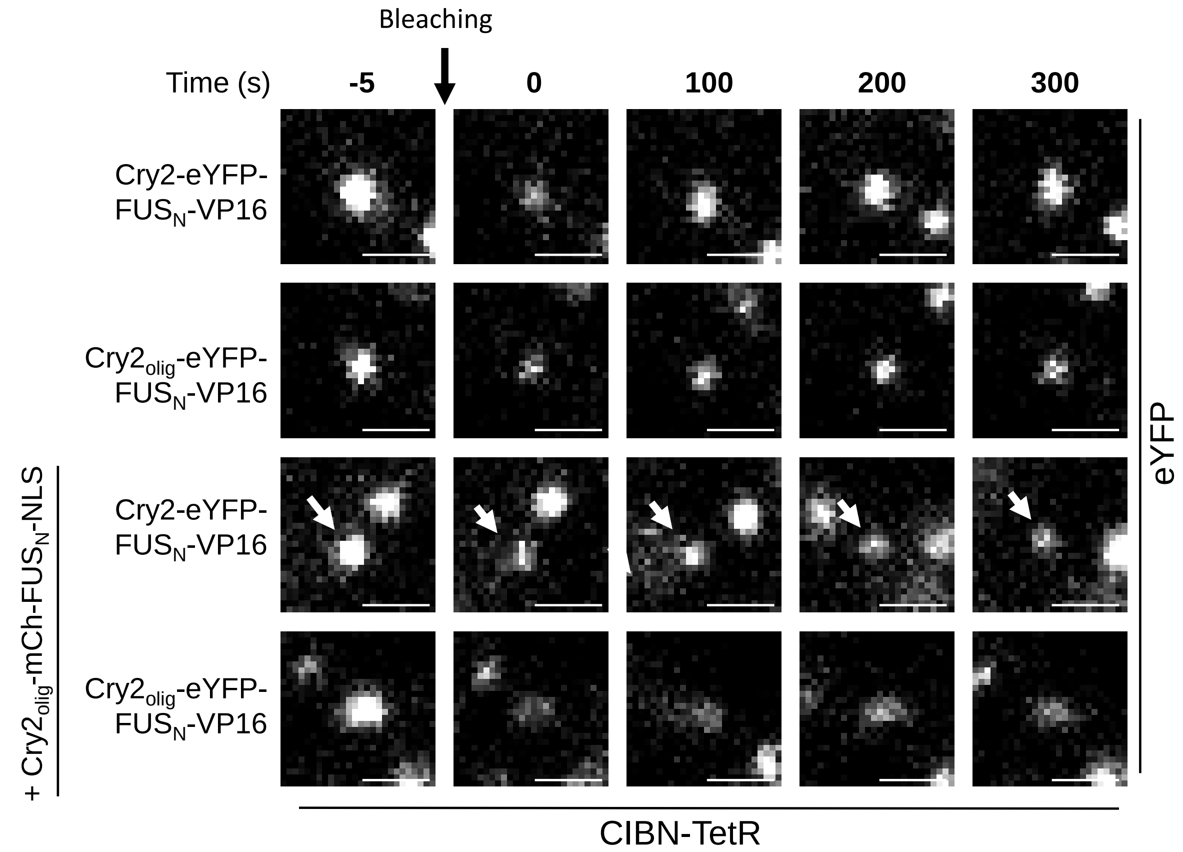

**Supplementary Figure 1: Bleaching and recovery of representative condensates**
CIBN-TetR + Cry2-eYFP-FUS$_N$-VP16 or + Cry2$_{olig}$-eYFP-FUS$_N$-VP16 constructs were transfected into HEK-293T cells either without or with the addition of Cry2$_{olig}$-mCh-FUS$_N$-NLS (1:2 ratio of plasmid amount). The cells were cultivated in the dark for 32 h prior to FRAP analysis. FRAP measurements were started after 10 min of blue light illumination (2.5 µmol m$^{-2}$ s$^{-1}$). Images show condensates before bleaching and selected time points of their recovery after bleaching. Scale bar = 1 µm.

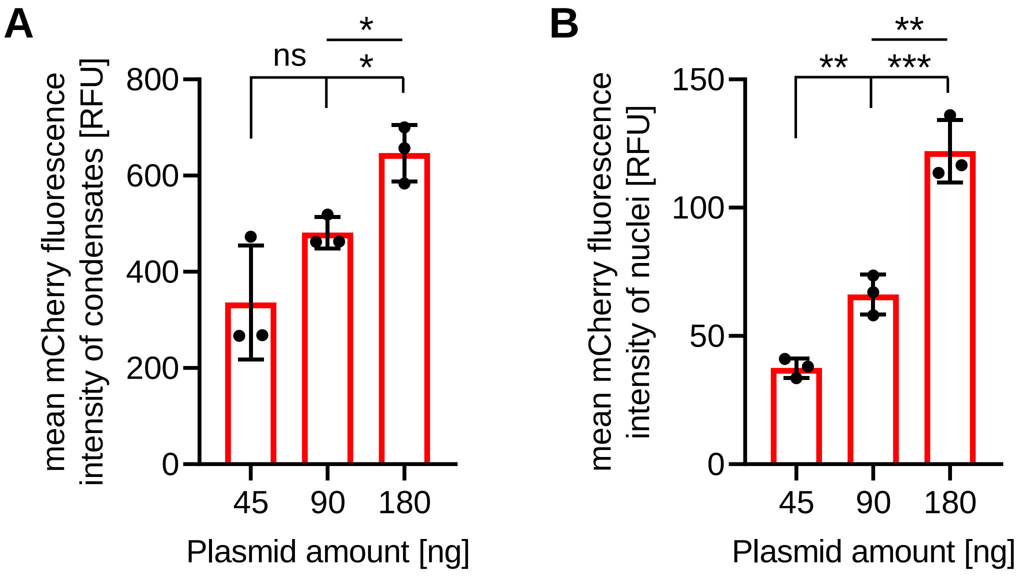

**Supplementary Figure 2: Quantitative analysis of the expression levels of Cry2$_{olig}$-mCh-FUS$_N$-NLS.**

Mean mCherry fluorescence intensity of condensates or nuclei. (**A, B**) HEK-293T cells were transfected with a tetO$_4$-based SEAP reporter together with CIBN-TetR, Cry2-eYFP-FUS$_N$-VP16 and increasing amounts of Cry2$_{olig}$-mCh-FUS$_N$-NLS (the indicated plasmid amounts correspond to the 1:1, 1:2 and 1:4 ratios in Figure 2). 8 h post transfection, cells were illuminated with blue light for 72 h (2.5 µmol m-$^2$ s-1) and afterwards analyzed by microscopy. For images, see Figure 2. (**A**) Mean mCherry fluorescence of the condensates. Single data points represent the median of the condensate population of one replicate, the bar represents the mean of n = 3 replicates ± SD. At least 251 condensates were analyzed per condition. (**B**) Mean mCherry fluorescence intensity of the nuclei. Background was subtracted, single data points represent the median of the nuclei intensities of one replicate, and the bar represents the mean of n = 3 replicates ± SD. At least 40 nuclei were analyzed per condition. Pairwise comparisons were performed using a Student's t.test (* = P ≤ 0.05; ** = P ≤ 0.01; *** = P ≤ 0.001).

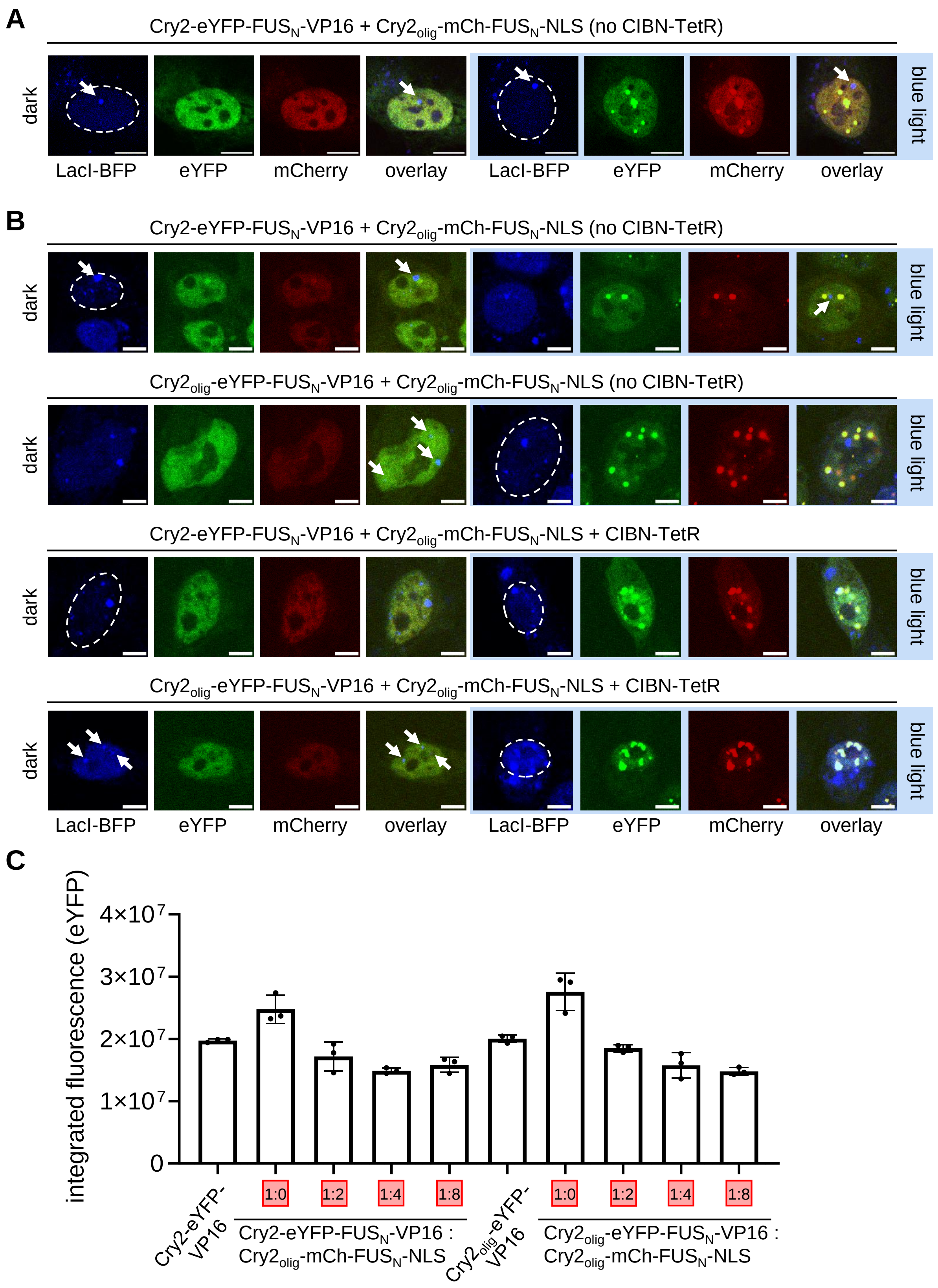

**Supplementary Figure 3: Controls for colocalization of transcription factor condensates with the target promoter and of eYFP expression levels**

(**A**) Control for colocalization of transcription factor condensates with the target promoter. U2-OS cells harboring a genomic locus with 256 lacO and 96 tetO repeats were transfected with constructs for LacI-BFP, Cry2-eYFP-FUS$_N$-VP16, and Cry2$_{olig}$-mCh-FUS$_N$-NLS. The cells were cultivated for 8 h, then illuminated for 24 h with blue light (2.5 µmol $m^{-2}$ $s^{-1}$) and subjected to microscopy analysis. Scale bar = 10 µm. (**B**) Colocalization of transcription factor condensates with the reporter plasmid. HEK-293T cells were transfected with the indicated constructs, LacI-BFP, and a reporter plasmid harboring 256 lacO and 6 tetO repeats. The cells were cultivated for 8 h, then illuminated for 24 h with blue light (5 µmol $m^{-2}$ $s^{-1}$), and subjected to microscopy analysis. Scale bar = 5 µm. (**C**) Integrated eYFP expression levels. The indicated constructs were transfected into HEK-293T cells together with constructs for DNA-binding (CIBN-TetR) and a $tetO_4$-based SEAP reporter. The numbers in the red boxes indicate the approximate plasmid ratio of the VP16 to mCherry-containing constructs. Cells were cultivated for 72 h in the dark prior determination of eYFP fluorescence via flow cytometry. Singlets were gated and eYFP expression levels were integrated. Data points represent the integrated eYFP expression levels ± SD (n = 3).

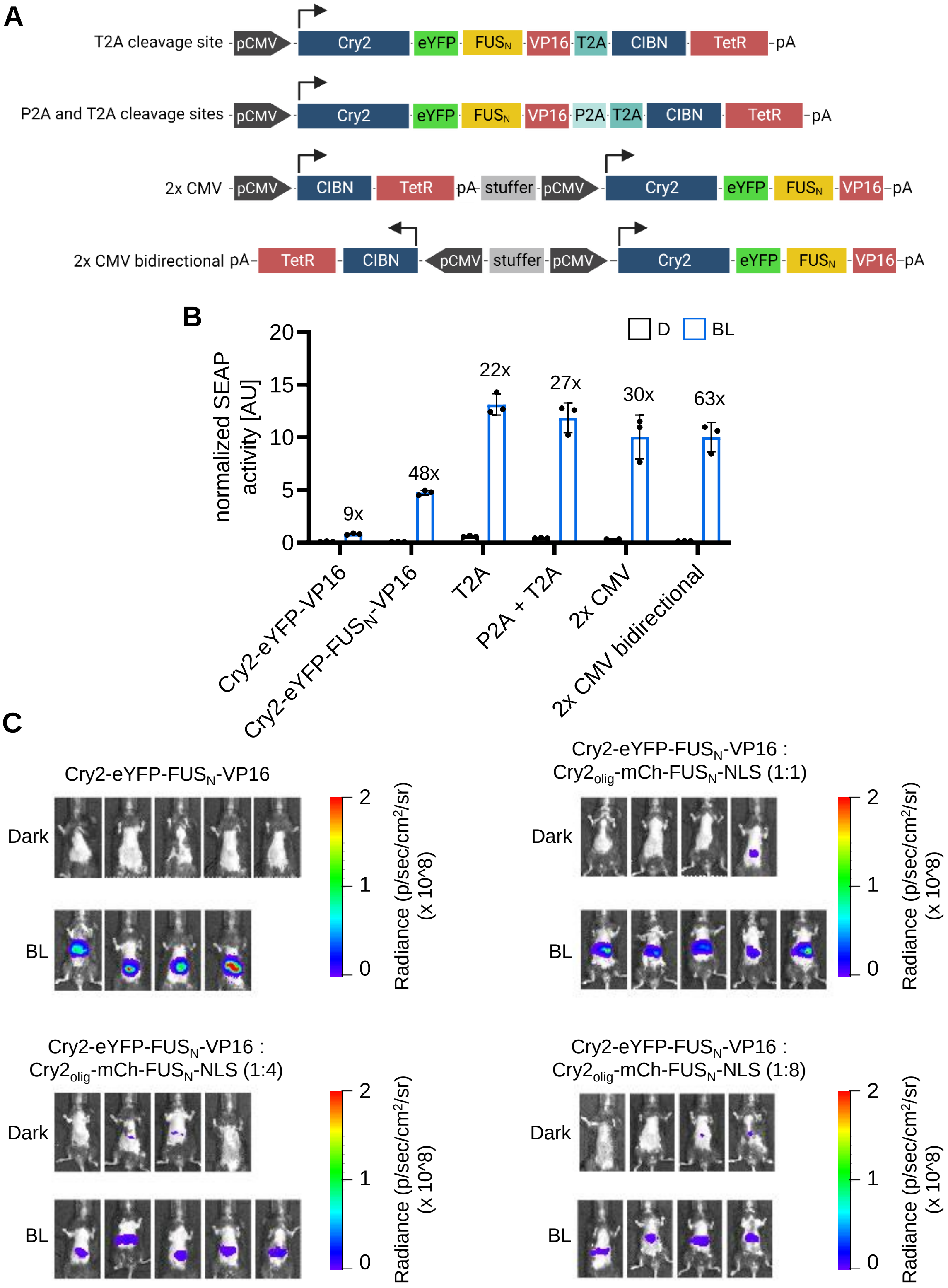
A
T2A cleavage site
pCMV
Cry2
eYFP
FUSN
VP16
T2A
CIBN
TetR
pA
P2A and T2A cleavage sites
P2A
2x CMV
stuffer
2x CMV bidirectional
B
normalized SEAP activity [AU]
D
BL
9x
48x
22x
27x
30x
63x
Cry2-eYFP-VP16
Cry2-eYFP-FUSN-VP16
T2A
P2A + T2A
2x CMV
2x CMV bidirectional
C
Cry2-eYFP-FUSN-VP16
Cry2-eYFP-FUSN-VP16 : Cry2olig-mCh-FUSN-NLS (1:1)
Cry2-eYFP-FUSN-VP16 : Cry2olig-mCh-FUSN-NLS (1:4)
Cry2-eYFP-FUSN-VP16 : Cry2olig-mCh-FUSN-NLS (1:8)
Dark
BL
Radiance (p/sec/cm²/sr) (x 10^8)

**Supplementary Figure 4: Design and testing of genetically compact expression vectors for mouse experiments**

(**A**) Design of vectors that express both Cry2-eYFP-FUS$_N$-VP16 and CIBN-TetR, separated either by posttranslational cleavage sites (T2A or a combination of T2A and P2A) or under the control of two CMV promoters. Either consecutively or bidirectional. (**B**) HEK-293T cells were transfected with a tetO$_4$-SEAP reporter and either Cry2-eYFP-FUS$_N$-VP16 and CIBN-TetR from two different plasmids or with constructs that express both components (as designed in (A)). 8 h after transfection, cells were either illuminated with blue light (2.5 µmol $m^{-2}$ $s^{-1}$) or kept in darkness for 72 h. SEAP activity was measured and additionally, cells were analyzed by flow cytometry to account for differences in expression levels of different constructs. SEAP values were normalized with the integrated eYFP fluorescence values from the dark samples. Means and single values ± SD are shown (n = 2-3). (**C**) Bioluminescence measurement images of mice that were hydrodynamically injected with the bidirectional vector encoding for CIBN-TetR and Cry2-eYFP-FUS$_N$-VP16, a tetO$_7$-based firefly luciferase reporter and the indicated ratios of Cry2$_{olig}$-mCh-FUS$_N$-NLS. Eight hours after plasmid injection, the mice were exposed to blue light pulses (460 nm, 10 mW $cm^{-2}$, 2 min on, 2 min off alternating) for 11 h. Subsequently, luciferin was injected intraperitoneally and bioluminescence images were acquired.

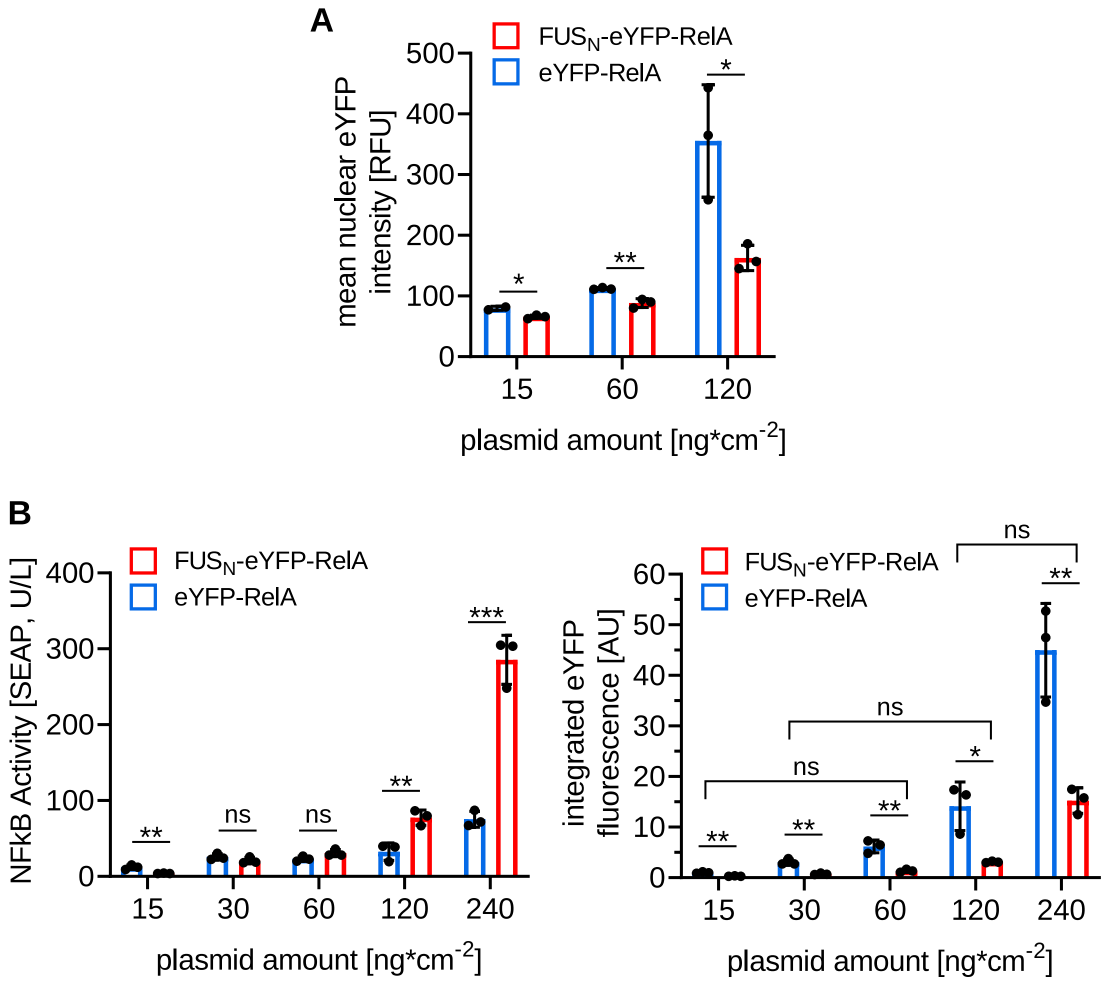

**Supplementary Figure 5: eYFP intensities of nuclei and extended titration of FUS$_N$-eYFP-RelA or eYFP-RelA**

(**A**) Quantitative analysis of the mean eYFP intensities of HEK-293T nuclei when transfected with FUS$_N$-eYFP-RelA or eYFP-RelA. At least 56 nuclei per condition were analyzed. Single data points represent the median of each replicate. Bars represent means ± SD (n = 2-3). **(B)** Titration of FUS$_N$-eYFP-RelA or eYFP-RelA. HEK-293T cells were transfected with the indicated amount of FUS$_N$-eYFP-RelA or eYFP-RelA expression vector and an NF-κB-responsive SEAP reporter. 32 h later, SEAP activity was measured and eYFP expression levels were determined by FACS. (left) mean ± SD and individual values are shown, n = 3. (right) mean ± SD and individual values are shown as fold change to the eYFP-RelA 15 ng*cm$^{-2}$ condition. The three conditions with non-significant differences in integrated eYFP levels (Arbitrary Units 1 (low), 3 (middle), 15 (high)) were used for the pairwise comparisons of SEAP production values in Figure 4C. *P* values were calculated using a Student‘s t.test: ns > 0.05; * = $P \leq 0.05$; ** = $P \leq 0.01$; *** = $P \leq 0.001$.

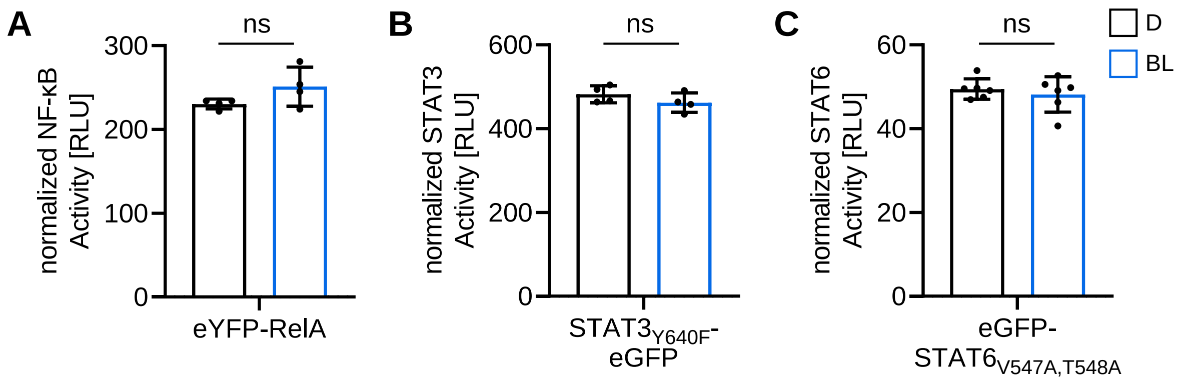

**Supplementary Figure 6: Control measurements for the influence of blue light on transcription factor activity**

(**A**) Influence of blue-light illumination on eYFP-RelA induced NF-κB activity. HEK-293T cells were transfected with eYFP-RelA, an NF-κB-responsive firefly luciferase reporter and a constitutive renilla luciferase reporter. 12 h post-transfection, cells were either kept in the dark or under blue-light illumination (5 µmol $m^{-2}$ $s^{-1}$) for 40 h prior to quantification of the luciferase activity. Values were normalized to constitutively expressed renilla luciferase activities. Means ± SD and single values are plotted; n = 4. (**B, C**) Impact of blue-light illumination on $STAT3_{Y640F}$-eGFP or eGFP-$STAT6_{V547A,T548A}$ activity. Cells were transfected with the indicated transcription factor, a STAT3 or STAT6-responsive firefly luciferase reporter, and a constitutive renilla reporter. 8 h after transfection, cells were stimulated with 5 ng/mL IL-6 for $STAT3_{Y640F}$ activation or 10 ng/mL IL-4 for $STAT6_{V547A,T548A}$ activation and then kept in the dark (D) or under blue light (BL) illumination (5 µmol $m^{-2}$ $s^{-1}$) for 24 h. Luciferase activities were quantified and firefly luciferase activity was normalized to renilla luciferase activity. Means ± SD and single values are plotted (n = 4 or 6). Pairwise comparisons were performed using a Student's t.test (ns = $P > 0.05$; * = $P \leq 0.05$).

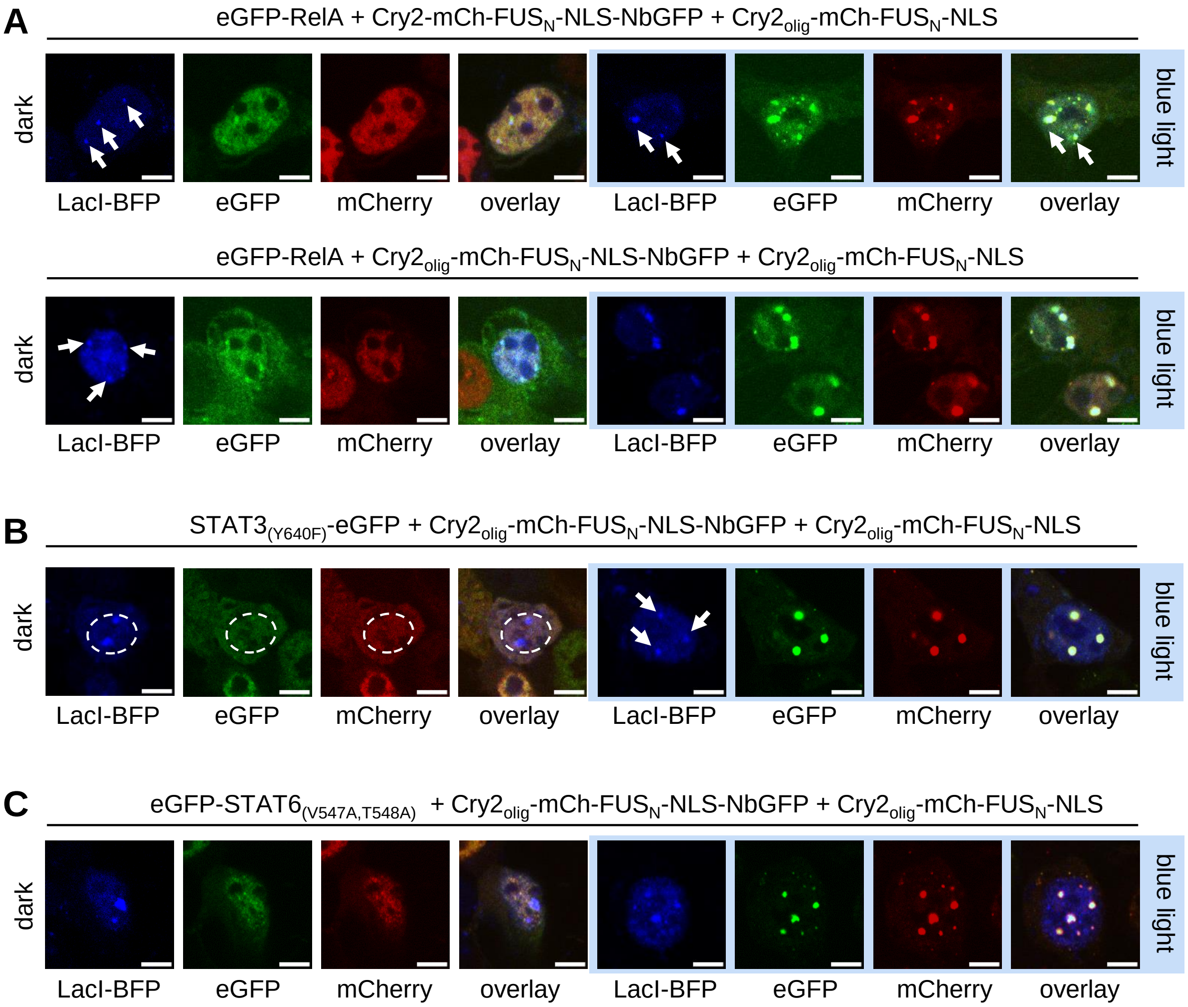

**Supplementary Figure 7: Colocalization of transcription factor condensates with reporter plasmids**

HEK-293T were transfected with the indicated constructs, LacI-BFP and a reporter plasmid containing an array of 256 lacO repeats and (**A**) an NF-κB response element, (**B**) a STAT3 response element or (**C**) a STAT6 response element. 8 h after transfection cells expressing $STAT3_{(Y640F)}$ or $STAT6_{(V547A,T548A)}$ were stimulated with 5 ng/mL IL-6 or 10 ng/mL IL-4, respectively. Afterwards, they were either kept in darkness or illuminated with blue light (5 µmol $m^{-2}$ $s^{-1}$) for 24 h, before fixation and analysis via microscopy. Please note: Minimum and maximum pixel values were adjusted differently between dark and blue light samples, to improve visibility. Scale bar = 5 µm.

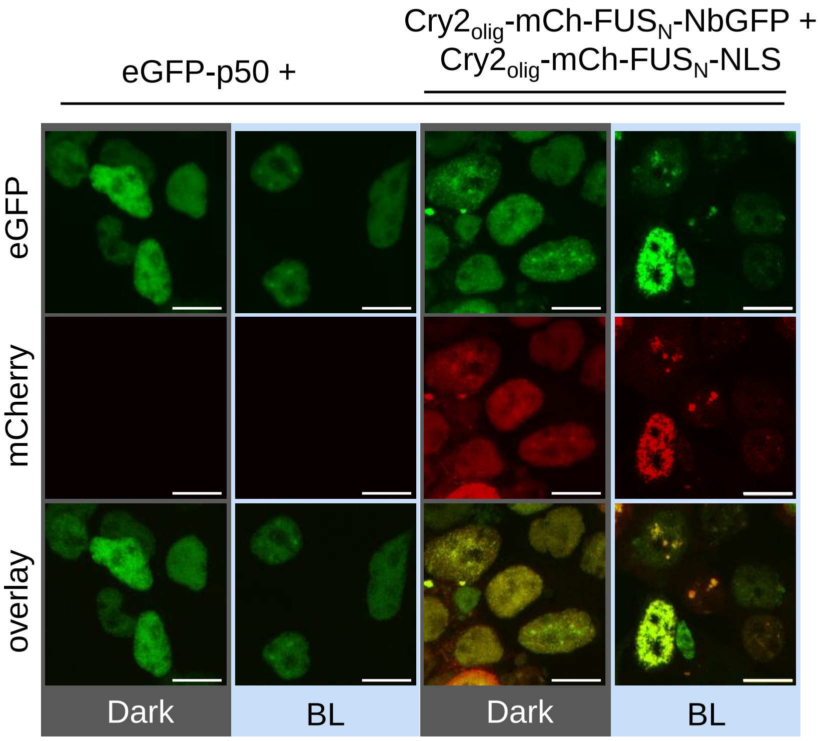

**Supplementary Figure 8: Microscopical analysis of the recruitment of eGFP-tagged p50 into condensates**

HEK293-T cells were transfected with the indicated constructs, an NF-κB-responsive firefly luciferase expression vector as well as a constitutive renilla luciferase construct. 8 h after transfection, cells were stimulated with 20 ng/mL TNF-α and then kept in the dark (D) or under blue light (BL) illumination (5 µmol $m^{-2}$ $s^{-1}$) for 24 h prior to microscopy analysis. Scale bar = 10 µm.

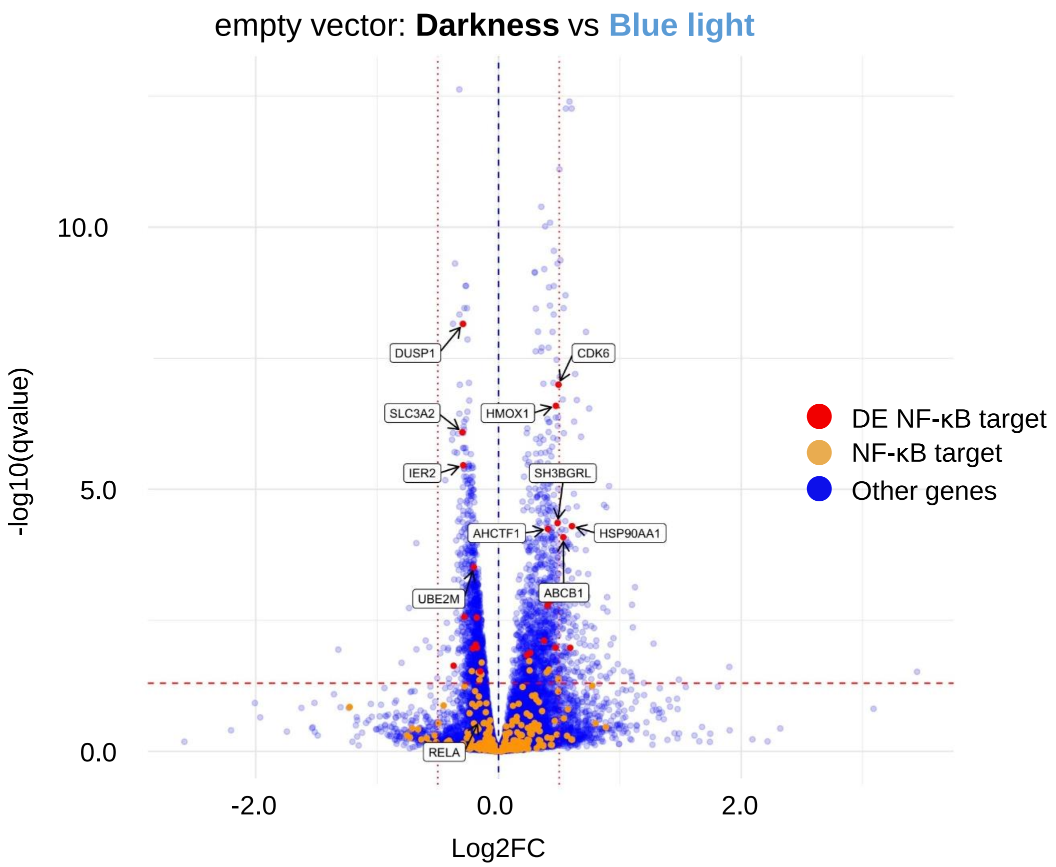
empty vector: Darkness vs Blue light
-log10(qvalue)
10.0
5.0
0.0
-2.0
0.0
2.0
Log2FC
DUSP1
CDK6
SLC3A2
HMOX1
IER2
SH3BGRL
AHCTF1
HSP90AA1
UBE2M
ABCB1
RELA
DE NF-κB target
NF-κB target
Other genes

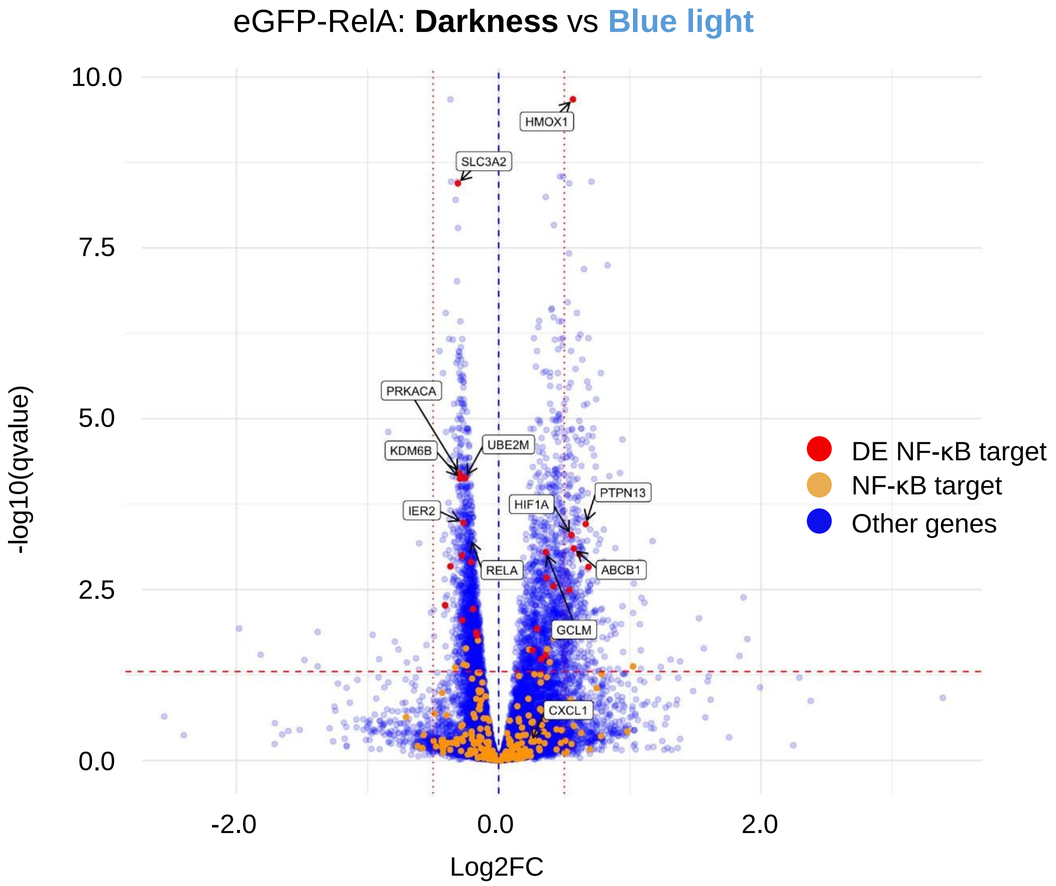
eGFP-RelA: Darkness vs Blue light
-log10(qvalue)
10.0
7.5
5.0
2.5
0.0
-2.0
0.0
2.0
Log2FC
HMOX1
SLC3A2
PRKACA
KDM6B
UBE2M
IER2
HIF1A
PTPN13
RELA
ABCB1
GCLM
CXCL1
DE NF-κB target
NF-κB target
Other genes

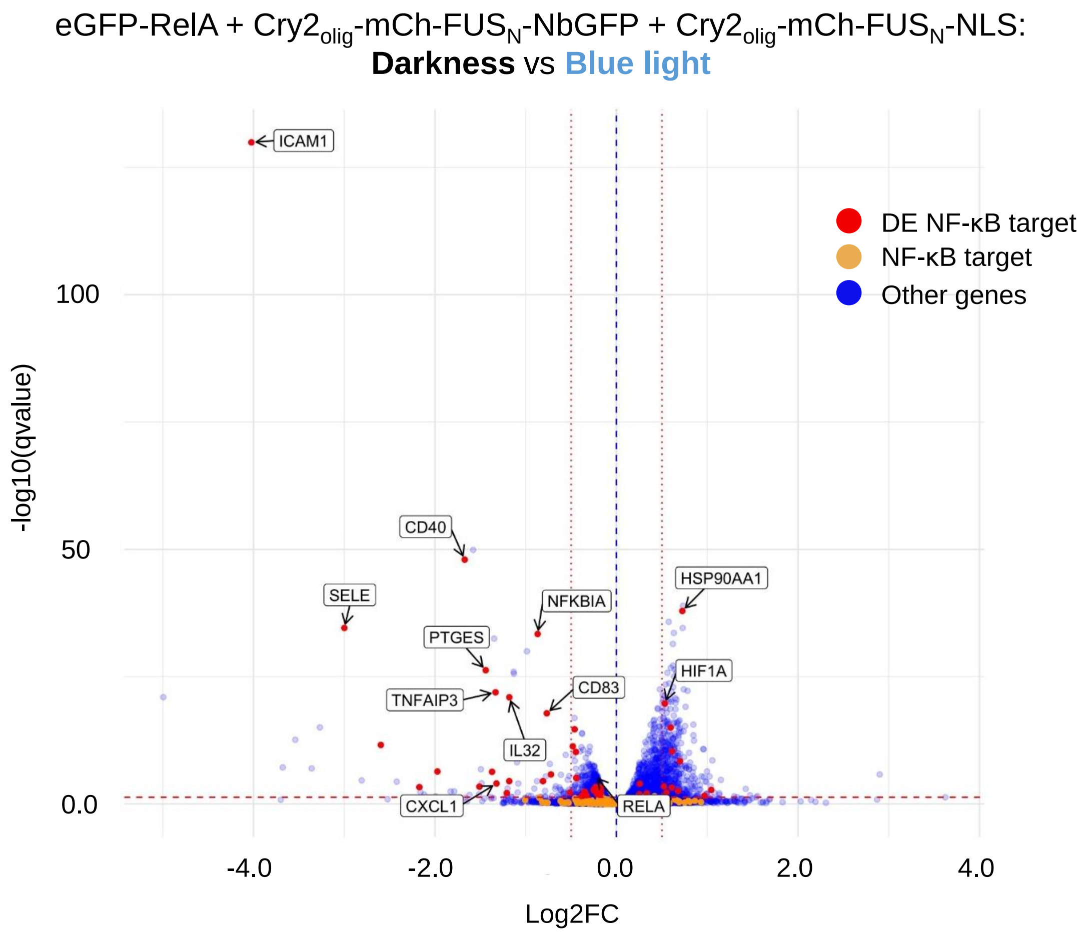

**Supplementary Figure 9: Deregulated genes between darkness and blue light.**
HEK-293T cells were transfected with the indicated expression vectors and an NF-κB-responsive SEAP reporter. 8 h after transfection, cells were either kept in the dark (D) or under blue light illumination (BL, 5 µmol m-² s-1) for 24 h prior to RNA extraction. Total RNA samples were subjected to RNAseq. Volcano plots show the comparison between darkness and blue-light conditions for the conditions indicated above each plot. Datasets of triplicates were pooled and genes with q-values ≤ 0.05 and | log2FC | > 0.5 were considered significantly deregulated. The orange points (NF-κB target genes) above the significance threshold are considered false positives since they are reported by the Wald Test as significantly differentially expressed but not by the Likelihood Ratio Test.

**Table S1: Numbers of deregulated genes, determination of the expression threshold, and filtering for NF-κB target genes**

The total number of statistically significantly deregulated (DE) genes with both q-values < 0.05 identified by Sleuth in the condensate condition reduced from 2313 to 932 after | log2FC | > 0.25 and to 143 after | log2FC | > 0.5 thresholds were applied, respectively. This corresponds to reduction factors of ~3 and ~16. In comparison, the deregulated NF-kB target genes in the condensate condition reduced from 51 to 36 after | log2FC | > 0.25 and to 21 after | log2FC | > 0.5 thresholds were applied, namely by factors of ~2 and ~3, respectively. By comparing the factors of reduction, we decided on the threshold 0.5 since it filtered out genes exhibiting un-specific effects without losing a high number of deregulated NF-kB target genes.

| | All Genes | | | All Genes, I log2FC I > 0.25 | | | All Genes, I log2FC I > 0.5 | | | NF-κB target genes | | | NF-κB target genes, I log2FC I > 0.25 | | | NF-κB target genes, I log2FC I > 0.5 | | |
|---|---|---|---|---|---|---|---|---|---|---|---|---|---|---|---|---|---|---|
| Comparison | Number of deregulated genes | | | Number of deregulated genes | | | Number of deregulated genes | | | Number of deregulated genes | | | Number of deregulated genes | | | Number of deregulated genes | | |
| Blue light: empty vector vs eGFP-RelA | 76 | upreg. | 65 | 71 | upreg. | 62 | 51 | upreg. | 49 | 26 | upreg. | 26 | 25 | upreg. | 25 | 21 | upreg. | 21 |
| | | downreg. | 11 | | downreg. | 9 | | downreg. | 2 | | downreg. | 0 | | downreg. | 0 | | downreg. | 0 |
| Darkness: empty vector vs eGFP-RelA | 66 | upreg. | 56 | 61 | upreg. | 53 | 44 | upreg. | 42 | 24 | upreg. | 24 | 24 | upreg. | 24 | 20 | upreg. | 20 |
| | | downreg. | 10 | | downreg. | 8 | | downreg. | 2 | | downreg. | 0 | | downreg. | 0 | | downreg. | 0 |
| Darkness vs. Blue light: eGFP-RelA + $Cry2_{olig}$-mCh-$FUS_N$-NLS-NbGFP + $Cry2_{olig}$-mCh-$FUS_N$-NLS | 2312 | upreg. | 1291 | 932 | upreg. | 801 | 143 | upreg. | 88 | 51 | upreg. | 14 | 36 | upreg. | 11 | 21 | upreg. | 3 |
| | | downreg. | 1021 | | downreg. | 131 | | downreg. | 55 | | downreg. | 37 | | downreg. | 25 | | downreg. | 18 |
| Darkness vs. Blue light: empty vector | 1678 | upreg. | 828 | 479 | upreg. | 462 | 26 | upreg. | 24 | 24 | upreg. | 13 | 12 | upreg. | 11 | 0 | upreg. | 0 |
| | | downreg. | 850 | | downreg. | 17 | | downreg. | 2 | | downreg. | 11 | | downreg. | 1 | | downreg. | 0 |
| Darkness vs. Blue light: eGFP-RelA | 2170 | upreg. | 1093 | 741 | upreg. | 705 | 56 | upreg. | 51 | 28 | upreg. | 15 | 12 | upreg. | 10 | 0 | upreg. | 0 |
| | | downreg. | 1077 | | downreg. | 36 | | downreg. | 5 | | downreg. | 13 | | downreg. | 2 | | downreg. | 0 |

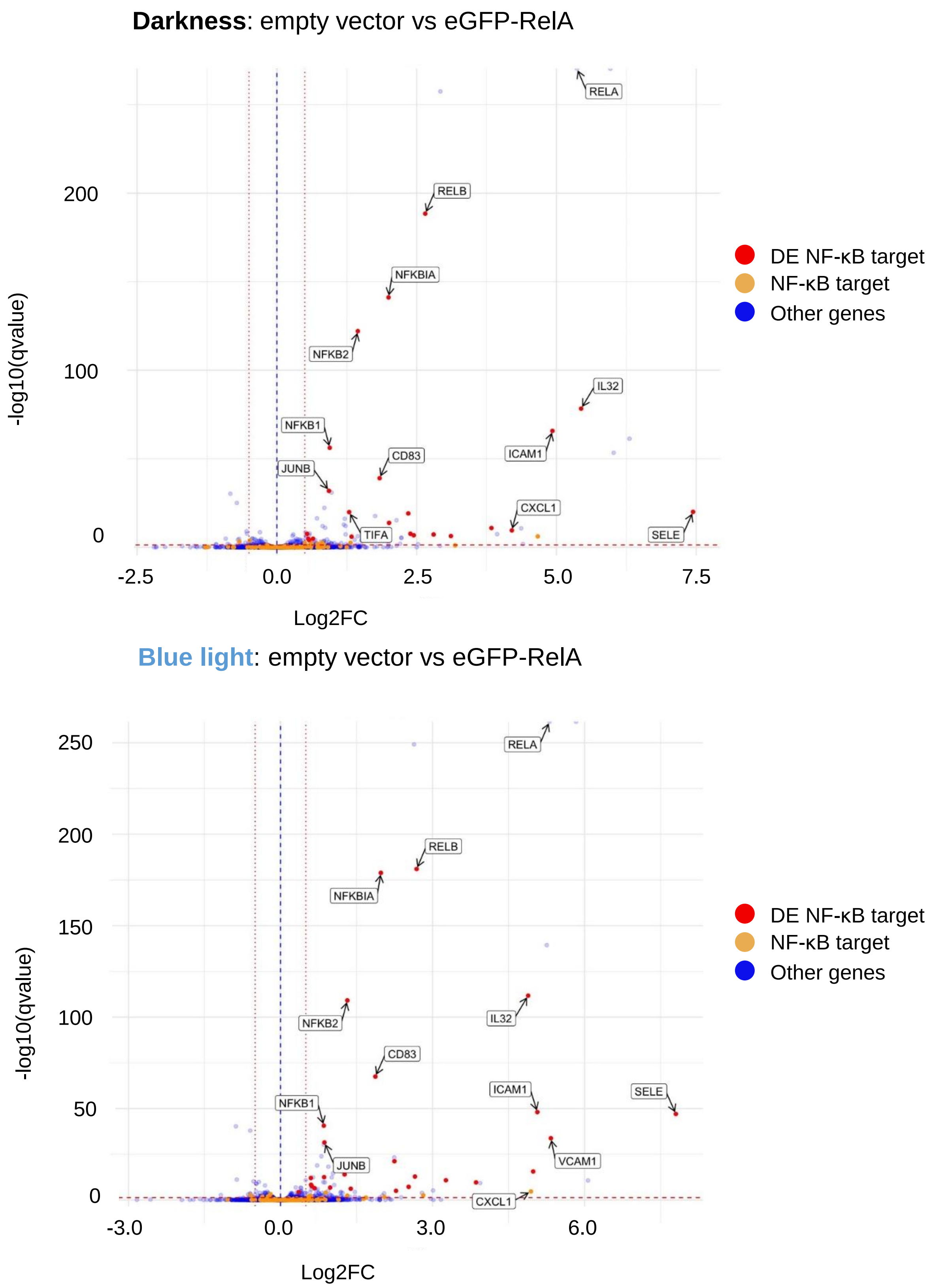

**Supplementary Figure 10**: **Identification of deregulated genes between empty vector and eGFP-RelA**

HEK-293T cells were treated as described in Figure S8. Volcano plots show the comparison between samples expressing empty vector vs eGFP-RelA. Datasets of triplicates were pooled and genes with q-values $\leq 0.05$ and $|\log2FC| > 0.5$ were considered significantly deregulated.

**NF-κB target genes | log2FC | > 0.5**

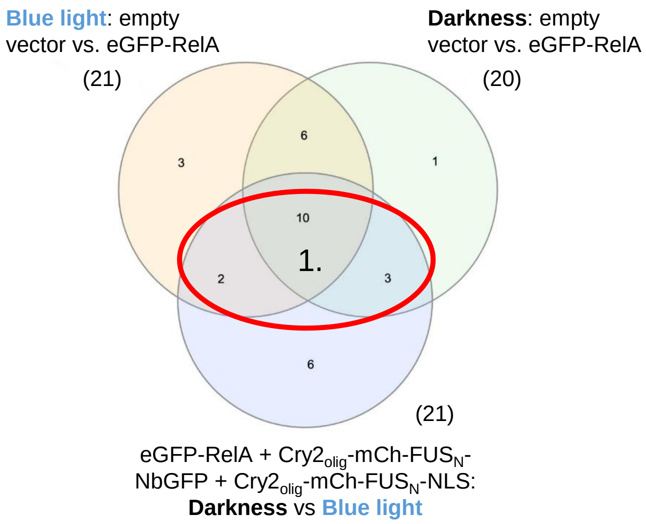

**1. Genes included in the heatmap (Figure 7)**

| target_ID | status* | qval_WT * | qval_LRT * |
|---|---|---|---|
| CD83 | - | 1.57E-18 | 0.00205 |
| CXCL1 | - | 9.60E-05 | 0.01212 |
| CXCL2 | - | 0.007171544 | 0.02512 |
| CXCL3 | - | 4.85E-07 | 0.00579 |
| CXCL8 | - | 4.22E-07 | 0.00436 |
| ICAM1 | - | 1.25E-130 | 0.00014 |
| IL32 | - | 1.13E-21 | 0.00245 |
| IRF1 | - | 3.28E-05 | 0.00941 |
| NFKBIA | - | 4.21E-34 | 0.00192 |
| PTGES | - | 5.46E-27 | 0.00186 |
| SELE | - | 2.75E-35 | 0.00148 |
| STAT5A | - | 2.57E-12 | 0.00257 |
| TIFA | - | 1.60E-06 | 0.00955 |
| TNF | - | 3.28E-05 | 0.00880 |
| TNFAIP3 | - | 1.19E-22 | 0.00205 |

* for eGFP-RelA + $Cry2_{olig}$-mCh-$FUS_N$-NbGFP + $Cry2_{olig}$-mCh-$FUS_N$-NLS: **Darkness** vs **Blue light**

**Supplementary Figure 11: Selection of the specifically deregulated genes and outlier analysis**

VENN diagram to determine the list of genes that are upregulated by eGFP-RelA and significantly deregulated by the condensate condition (eGFP-RelA + $Cry2_{olig}$-mCh-$FUS_N$-NbGFP + $Cry2_{olig}$-mCh-$FUS_N$-NLS) after blue-light illumination. The status "-" indicates downregulation, q-values (qval) were determined with the Wald Test (WT) and the Likelihood Ratio Test (LRT). If both q-values ≤ 0.05 and | log2FC | > 0.5, the deregulation is considered true positive.

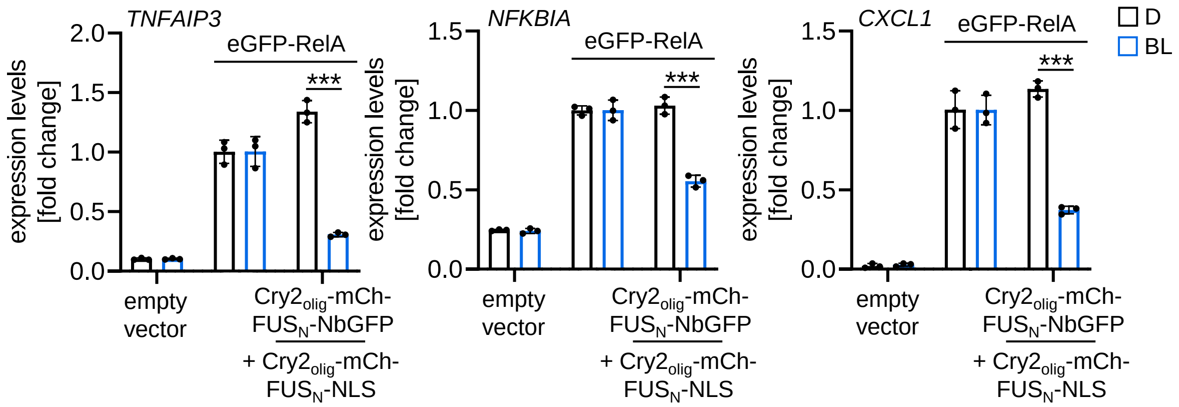

**Supplementary Figure 12: Impact of recruitment of RelA into condensates on endogenous gene expression.**
HEK-293T cells were transfected with the indicated expression vectors and an NF-κB-responsive SEAP reporter. 8 h after transfection, cells were either kept in the dark (D) or under blue light illumination (BL, 5 µmol $m^{-2}$ $s^{-1}$) for 24 h prior to RNA extraction, reverse transcription, and qPCR analysis. Data was normalized to the housekeeping gene GUS. Mean $2^{\wedge -\Delta\Delta Ct}$ values ± SD are plotted as fold changes to the respective eGFP-RelA-only conditions, n = 3. *P* values were calculated using a Student's t.test on the ΔΔCt values (*** = P ≤ 0.001).

**Table S2: Plasmids used in this study**

| Category | Name | Description | Backbone |
|---|---|---|---|
| Reporters | pNS304 | $tetO_4$-SEAP reporter | pKM001 |
| | pAF101 | $tetO_7$-Firefly luciferase reporter | pKM001 |
| | pAF504 | 3x NF-κB-RE-SEAP reporter | pMF111 |
| | NF-κB-Firefly luciferase | 3x NF-κB-RE-Firefly luciferase reporter | Ref. 45 |
| | pAF133 | 3x STAT3-RE-Firefly luciferase reporter | tdTomato-N1 |
| | pAF312 | STAT6-RE-Firefly luciferase reporter | pHW040 |
| | Renilla lucif-erase | pCMV-Renilla luciferase | pRL-CMV |
| | TK-Renilla lu-ciferase | pTK-Renilla luciferase | pRL-TK |
| | pNS053 | $lacO_{256}$-$tetO_6$-$P_{hCMVmin}$-SEAP | pKM001 |
| | pAF186 | $lacO_{256}$-NF-κB-$RE_3$-$tetO_6$-$P_{hCMVmin}$-SEAP | pNS053 |
| | pAF187 | $lacO_{256}$-STAT3-RE-$tetO_6$-$P_{hCMVmin}$-SEAP | pNS053 |
| | pAF188 | $lacO_{256}$-STAT6-RE-$tetO_6$-$P_{hCMVmin}$-SEAP | pNS053 |
| General | pAF057 | empty vector | pEGFP-C3 |
| | pAF082 | $Cry2_{olig}$-mCh-$FUS_N$-NLS | pEGFP-C3 |
| | pAF525 | LacI-BFP | pN2 |
| Syn-theticTF | pNS1000 | CIBN-TetR | pEGFP-C3 |
| | pNS1001 | Cry2-eYFP-VP16 | pEGFP-C3 |
| | pNS026 | Cry2-eYFP-$FUS_N$-VP16 | pEGFP-C3 |
| | pAF500 | $Cry2_{olig}$-eYFP-$FUS_N$-VP16 | pEGFP-C3 |
| | pAF501 | $Cry2_{olig}$-eYFP-VP16 | pEGFP-C3 |
| | pAF514 | pCMV-CIBN-TetR-pCMV-Cry2-eYFP-$FUS_N$-VP16 | pEGFP-C3 |
| | pAF515 | TetR-CIBN-pCMV-stuffer-pCMV-Cry2-eYFP-$FUS_N$-VP16 (bidirectional) | pEGFP-C3 |
| | pAF517 | Cry2-eYFP-$FUS_N$-VP16-T2A-CIBN-TetR | pEGFP-C3 |
| | pAF518 | Cry2-eYFP-$FUS_N$-VP16-P2A-T2A-CIBN-TetR | pEGFP-C3 |
| | pAF519 | TetR-CIBN-pCMV-stuffer-pCMV-Cry2-eYFP-VP16 (bidirectional) | pEGFP-C3 |
| NF-κB | pAF521 | eYFP-RelA | pEGFP-C3 |
| | pAF522 | $FUS_N$-eYFP-RelA | pEGFP-C3 |
| | pAF510 | Cry2-eYFP-$FUS_N$-RelA | pEGFP-C3 |
| | pAF513 | $Cry2_{olig}$-eYFP-$FUS_N$-RelA | pEGFP-C3 |
| | pAF149 | eGFP-RelA | pEGFP-C3 |
| | pAF301 | $Cry2_{olig}$-mCh-$FUS_N$-NLS-NbGFP | pEGFP-C3 |
| | pAF353 | Cry2-mCh-$FUS_N$-NLS-RelA | pEGFP-C3 |
| | eGFP-p50 | eGFP-p50 | pEGFP-C3 |
| STAT | pAF167 | STAT3(Y640F)-eGFP | pEGFP-C3 |
| | pAF326 | eGFP-STAT6(V547A,T548A) | pEGFP-C3 |

**Table S3: Transfection conditions of each experiment**

| Figure | Format | Condition | Plasmidname and DNA amount |
|---|---|---|---|
| Figure 1B, S1 | 32 mm Ibidi | CIBN-TetR + Cry2/Cry2$_{olig}$-eYFP-FUS$_N$-VP16 | pNS1000 (92 ng), pNS026/pAF500 (92 ng), pNS304 (2300 ng), pAF057 (966 ng) |
| | | CIBN-TetR + Cry2/Cry2$_{olig}$-eYFP-FUS$_N$-VP16 + Cry2$_{olig}$-mCh-FUS$_N$-NLS | pNS1000 (92 ng), pNS026/pAF500 (92 ng), pNS304 (2300 ng), pAF057 (769 ng), pAF082 (197 ng) |
| Figure 2A,B, Fig. S2 | 24-well | CIBN-TetR + Cry2/Cry2$_{olig}$-eYFP-VP16 | pNS1000 (45 ng) + pNS1001/pAF501 (45 ng) + pNS304 (480 ng) + pAF057 (180 ng) |
| | | CIBN-TetR + Cry2/Cry2$_{olig}$-eYFP-FUS$_N$-VP16 | pNS1000 (45 ng) + pNS026/pAF500 (45 ng) + pNS304 (480 ng) + pAF057 (180 ng) |
| | | CIBN-TetR + Cry2-eYFP-FUS$_N$-VP16 + Cry2$_{olig}$-mCh-FUS$_N$-NLS (1:1) | pNS1000 (45 ng) + pNS026 (45 ng) + pNS304 (480 ng) + pAF057 (135 ng) + pAF082 (45 ng) |
| | | CIBN-TetR + Cry2-eYFP-FUS$_N$-VP16 + Cry2$_{olig}$-mCh-FUS$_N$-NLS (1:2) | pNS1000 (45 ng) + pNS026 (45 ng) + pNS304 (480 ng) + pAF057 (90 ng) + pAF082 (90 ng) |
| | | CIBN-TetR + Cry2-eYFP-FUS$_N$-VP16 + Cry2$_{olig}$-mCh-FUS$_N$-NLS (1:4) | pNS1000 (45 ng) + pNS026 (45 ng) + pNS304 (480 ng) + pAF082 (180 ng) |
| Figure 3A, S3A | 24-well | LacI-BFP + Cry2-eYFP-FUS$_N$ -VP16 + Cry2$_{olig}$-mCh-FUS$_N$-NLS | pAF525 (300 ng) + pNS026 (150 ng) + pAF082 (150 ng) + pAF057 (150 ng) |
| | | LacI-BFP + CIBN-TetR + Cry2-eYFP-FUS$_N$-VP16 + Cry2$_{olig}$-mCh-FUS$_N$-NLS (1:1) | pAF525 (300 ng) + pNS1000 (150 ng) + pNS026 (150 ng) + pAF082 (150 ng) + pAF057 (150 ng) |
| Figure 3B, S3C | 96-well | CIBN-TetR + Cry2/Cry2$_{olig}$-eYFP-VP16 | pNS1000 (3.5 ng) + pNS1001/pAF501 (3.5 ng) + pNS304 (85 ng) + pAF057 (33 ng) |
| | | CIBN-TetR + Cry2/Cry2$_{olig}$-eYFP-FUS$_N$-VP16 | pNS1000 (3.5 ng) + pNS026/pAF500 (3.5 ng) + pNS304 (85 ng) + pAF057 (33 ng) |
| | | CIBN-TetR + Cry2/Cry2$_{olig}$-eYFP-FUS$_N$-VP16 + Cry2$_{olig}$-mCh-FUS$_N$-NLS (1:2) | pNS1000 (3.5 ng) + pNS026/pAF500 (3.5 ng) + pNS304 (85 ng) + pAF057 (25.5 ng) + pAF082 (7.5 ng) |
| | | CIBN-TetR + Cry2/Cry2$_{olig}$-eYFP-FUS$_N$-VP16 + Cry2$_{olig}$-mCh-FUS$_N$-NLS (1:4) | pNS1000 (3.5 ng) + pNS026/pAF500 (3.5 ng) + pNS304 (85 ng) + pAF057 (18 ng) + pAF082 (15 ng) |
| | | CIBN-TetR + Cry2/Cry2$_{olig}$-eYFP-FUS$_N$-VP16 + Cry2$_{olig}$-mCh-FUS$_N$-NLS (1:8) | pNS1000 (3.5 ng) + pNS026/pAF500 (3.5 ng) + pNS304 (85 ng) + pAF057 (3 ng) + pAF082 (30 ng) |
| | Mouse, *in vivo* | CIBN-TetR + Cry2-eYFP-FUS$_N$-VP16 (bidirectional) | pAF101 (240 µg) + pAF515 (10 µg) + pAF057 (80 µg) |

| | | | |
|---|---|---|---|
| Figure 3C, S4C | | CIBN-TetR + Cry2-eYFP-FUS$_{N}$-VP16 (bidirectional) + Cry2$_{olig}$-mCh-FUS$_{N}$-NLS 1:1 | pAF101 (240 µg) + pAF515 (10 µg) + pAF082 (10 µg) + pAF057 (70 µg) |
| | | CIBN-TetR + Cry2-eYFP-FUS$_{N}$-VP16 (bidirectional) + Cry2$_{olig}$-mCh-FUS$_{N}$-NLS 1:4 | pAF101 (240 µg) + pAF515 (10 µg) + pAF082 (40 µg) + pAF057 (40 µg) |
| | | CIBN-TetR + Cry2-eYFP-FUSN-VP16 (bidirectional) + Cry2$_{olig}$-mCh-FUS$_{N}$-NLS 1:8 | pAF101 (240 µg) + pAF515 (10 µg) + pAF082 (80 µg) |
| Figure 4A, B, S5A | 24-well | FUS$_{N}$ -eYFP-RelA titration (3 conditions) | pAF504 (300 ng) + pAF522 (30 - 120 - 240 ng) + pAF057 (420 - 330 - 210 ng) |
| | | eYFP-RelA titration (3 conditions) | pAF504 (300 ng) + pAF521 (30 - 120 - 240 ng) + pAF057 (420 - 330 - 210 ng) |
| Figure 4C, S5B | 96-well | FUS$_{N}$-eYFP-RelA titration (5 conditions) | pAF504 (45 ng) + pAF522 (5 - 10 - 20 - 40 - 80 ng) + pAF057 (75 - 70 - 60 – 40 - 0 ng) |
| | | eYFP-RelA titration (5 conditions) | AF504 (45 ng) + pAF521 (5 - 10 - 20 - 40 - 80 ng) + pAF057 (75 - 70 - 60 – 40 - 0 ng) |
| Figure 5A, B | 24-well | Cry2-eYFP-FUS$_{N}$-RelA + Cry2$_{olig}$-mCh-FUS$_{N}$-NLS | pAF504 (150 ng) + pTK-Renilla luciferase (120 ng) + pAF510 (240 ng) + pAF082 (120 ng) + pAF057 (120 ng) |
| | | Cry2$_{olig}$-eYFP-FUS$_{N}$-RelA + Cry2$_{olig}$-mCh-FUS$_{N}$-NLS | pAF504 (150 ng) + pTK-Renilla luciferase (120 ng) + pAF513 (240 ng) + pAF082 (120 ng) + pAF057 (120ng) |
| Figure 5C | 96-well | Cry2-eYFP-FUS$_{N}$-RelA + Cry2$_{olig}$-mCh-FUS$_{N}$-NLS | NF-κB-Firefly luciferase (25 ng) + pCMV-Renilla luciferase (20 ng) + pAF510 (40 ng) + pAF082 (20 ng) + pAF057 (20 ng) |
| | | Cry2$_{olig}$-eYFP-FUS$_{N}$-RelA + Cry2$_{olig}$-mCh-FUS$_{N}$-NLS | NF-κB-Firefly luciferase (25 ng) + pCMV-Renilla luciferase (20 ng) + pAF513 (40 ng) + pAF082 (20 ng) + pAF057 (20 ng) |
| Figure 6A | 24-well | eGFP-RelA | NF-κB-Firefly luciferase (150 ng) + pTK-Renilla luciferase (120 ng) + pAF149 (6 ng) + pAF057 (474 ng) |
| | | eGFP-RelA + Cry2-mCh-FUS$_{N}$-NbGFP + Cry2$_{olig}$-mCh-FUS$_{N}$-NLS | NF-κB-Firefly luciferase (150 ng) + pTK-Renilla luciferase (120 ng) + pAF149 (6 ng) + pAF353 (240 ng) + pAF082 (120 ng) + pAF057 (144 ng) |
| | | eGFP-RelA + Cry2$_{olig}$-mCh-FUS$_{N}$-NbGFP + Cry2$_{olig}$-mCh-FUS$_{N}$-NLS | NF-κB-Firefly luciferase (150 ng) + pTK-Renilla luciferase (120 ng) + pAF149 (6 ng) + pAF301 (240 ng) + pAF082 (120 ng) + pAF057 (144 ng) |
| Figure 6B | 96-well | reporters only | NF-κB-Firefly luciferase (25 ng) + pCMV-Renilla luciferase (20 ng) + pAF057 (80 ng) |

| | | | |
|---|---|---|---|
| | | eGFP-RelA | NF-κB-Firefly luciferase (25 ng) + pCMV-Renilla luciferase (20 ng) + pAF149 (1 ng) + pAF057 (79 ng) |
| | | eGFP-RelA + Cry2-mCh-$FUS_N$-NbGFP + $Cry2_{olig}$-mCh-$FUS_N$-NLS | NF-κB-Firefly luciferase (25 ng) + pCMV-Renilla luciferase (20 ng) + pAF149 (1 ng) + pAF353 (40 ng) + pAF082 (20 ng) + pAF057 (19 ng) |
| | | eGFP-RelA + $Cry2_{olig}$-mCh-$FUS_N$-NbGFP + $Cry2_{olig}$-mCh-$FUS_N$-NLS | NF-κB-Firefly luciferase (25 ng) + pCMV-Renilla luciferase (20 ng) + pAF149 (1 ng) + pAF301 (40 ng) + pAF082 (20 ng) + pAF057 (19 ng) |
| Figure 6C | 96-well | reporters only (-/+ IL-6) | pAF133 (30 ng) + pCMV-Renilla luciferase (20 ng) + pAF057 (75 ng) |
| | | $STAT3_{(Y640F)}$-eGFP + $Cry2_{olig}$-mCh-$FUS_N$-NbGFP + $Cry2_{olig}$-mCh-FUSN-NLS | pAF133 (30 ng) + pCMV-Renilla luciferase (20 ng) + pAF167 (10 ng) + pAF301 (40 ng) + pAF082 (20 ng) + pAF057 (5 ng) |
| Figure 6D | 96-well | eGFP-$STAT6_{(V547A,T548A)}$ (-/+ IL-4) | pAF312 (30 ng) + pTK-Renilla luciferase (20 ng) + pAF326 (10 ng) + pAF057 (65 ng) |
| | | eGFP-$STAT6_{(V547A,T548A)}$ + $Cry2_{olig}$-mCh-$FUS_N$-NbGFP + $Cry2_{olig}$-mCh-$FUS_N$-NLS | pAF312 (30 ng) + pTK-Renilla luciferase (20 ng) + pAF326 (10 ng) + pAF301 (40 ng) + pAF082 (25 ng) |
| Figure 6E | 96-well | reporters only (-/+ TNF-α) | NF-κB-Firefly luciferase (25 ng) + pCMV-Renilla luciferase (20 ng) + pAF057 (80 ng) |
| | | eGFP-p50 | NF-κB-Firefly luciferase (25 ng) + pCMV-Renilla luciferase (20 ng) + eGFP-p50 (5 ng) + pAF057 (75 ng) |
| | | eGFP-p50 + $Cry2_{olig}$-mCh-$FUS_N$-NbGFP + $Cry2_{olig}$-mCh-$FUS_N$-NLS | NF-κB-Firefly luciferase (25 ng) + pCMV-Renilla luciferase (20 ng) + eGFP-p50 (5 ng) + pAF301 (35 ng) + pAF082 (20 ng) + pAF057 (15 ng) |
| Figure 7, S9-12 | 6-well | reporter only | pAF504 (1200 ng) + pAF057 (2550 ng) |
| | | eGFP-RelA | pAF504 (1200 ng) + pAF149 (30 ng) + pAF057 (2520 ng) |
| | | eGFP-RelA + $Cry2_{olig}$-mCh-FUSN-NbGFP + $Cry2_{olig}$-mCh-$FUS_N$-NLS | pAF504 (1200 ng) + pAF149 (30 ng) + pAF301 (1200 ng) + pAF082 (600 ng) + pAF057 (720 ng) |
| Figure S3B | 24-well | LacI-BFP + Cry2-eYFP-FUSn-VP16 + $Cry2_{olig}$-mCh-FUSN-NLS | pAF525 (90 ng) + pNS026 (90 ng) + pAF082 (180 ng) + pAF057 (90 ng) + pNS053 (300 ng) |
| | | LacI-BFP + $Cry2_{olig}$-eYFP-$FUS_N$ -VP16 + $Cry2_{olig}$-mCh-$FUS_N$-NLS | pAF525 (90 ng) + pAF500 (90 ng) + pAF082 (180 ng) + pAF057 (90 ng) + pNS053 (300 ng) |
| | | LacI-BFP + CIBN-TetR + Cry2-eYFP- $FUS_N$ -VP16 + $Cry2_{olig}$-mCh-$FUS_N$-NLS | pAF525 (90 ng) + pNS1000 (90 ng) + pNS026 (90 ng) + pAF082 (180 ng) + pAF057 (90 ng) + pNS053 (300 ng) |

| | | | |
|---|---|---|---|
| | | LacI-BFP + CIBN-TetR + $Cry2_{olig}$-eYFP- $FUS_N$ -VP16 + $Cry2_{olig}$-mCh-$FUS_N$-NLS | pAF525 (90 ng) + pNS1000 (90 ng) + pAF500 (90 ng) + pAF082 (180 ng) + pAF057 (90 ng) + pNS053 (300 ng) |
| Figure S4A, B | 96-well | CIBN-TetR + Cry2-eYFP-VP16 | pNS304 (85 ng) + pNS1000 (3,5 ng) + pNS1001 (3,5 ng) + pAF057 (33 ng) |
| | | CIBN-TetR + Cry2-eYFP-$FUS_N$-VP16 | pNS304 (85 ng) + pNS1000 (3,5 ng) + pNS026 (3,5 ng) + pAF057 (33 ng) |
| | | CIBN-TetR + Cry2-eYFP-$FUS_N$-VP16 separated by T2A | pNS304 (85 ng) + pAF517 (3,5 ng) + pAF057 (36,5 ng) |
| | | CIBN-TetR + Cry2-eYFP-$FUS_N$-VP16 separated by T2A + P2A | pNS304 (85 ng) + pAF518 (3,5 ng) + pAF057 (36,5 ng) |
| | | CIBN-TetR + Cry2-eYFP-$FUS_N$-VP16 separated by 2x pCMV | pNS304 (85 ng) + pAF514 (3,5 ng) + pAF057 (36,5 ng) |
| | | CIBN-TetR + Cry2-eYFP-$FUS_N$-VP16 separated by 2x pCMV bidirectional | pNS304 (85 ng) + pAF515 (3,5 ng) + pAF057 (36,5 ng) |
| Figure S6A | 96 - well | eYFP-RelA | NF-κB-Firefly luciferase (25 ng) + pCMV-Renilla luciferase (20 ng) + pAF521 (10 ng) + pAF057 (70 ng) |
| Fig-ure. S6B | 96 - well | $STAT3_{(Y640F)}$-eGFP + IL-6 | pAF133 (30 ng) + pTK-Renilla luciferase (20 ng) + pAF167 (10 ng) + pAF057 (65 ng) |
| Figure S6C | 96 - well | eGFP-$STAT6_{(V547A,T548A)}$ + IL-4 | pAF312 (30 ng) + pTK-Renilla luciferase (20 ng) + pAF326 (10 ng) + pAF057 (65 ng) |
| Figure S7 | 24-well | LacI-BFP + eGFP-RelA + Cry2-mCh-$FUS_N$-NbGFP + $Cry2_{olig}$-mCh-$FUS_N$-NLS | pAF525 (80 ng) + pAF186 (260 ng) + pAF149 (3 ng) + pAF353 (120 ng) + pAF082 (60 ng) + pAF057 (227 ng) |
| | | LacI-BFP + eGFP-RelA + $Cry2_{olig}$-mCh-$FUS_N$-NbGFP + $Cry2_{olig}$-mCh-$FUS_N$-NLS | pAF525 (80 ng) + pAF186 (260 ng) + pAF149 (3 ng) + pAF301 (120 ng) + pAF082 (60 ng) + pAF057 (227 ng) |
| | | LacI-BFP + $STAT3_{(Y640F)}$-eGFP + $Cry2_{olig}$-mCh-$FUS_N$-NbGFP + $Cry2_{olig}$-mCh-$FUS_N$-NLS | pAF525 (80 ng) + pAF187 (260 ng) + pAF167 (30 ng) + pAF301 (120 ng) + pAF082 (60 ng) + pAF057 (200 ng) |
| | | LacI-BFP + eGFP-$STAT6_{(V547A,T548A)}$ + $Cry2_{olig}$-mCh-$FUS_N$-NbGFP + $Cry2_{olig}$-mCh-$FUS_N$-NLS | pAF525 (80 ng) + pAF188 (260 ng) + pAF326 (30 ng) + pAF301 (120 ng) + pAF082 (75 ng) + pAF057 (185 ng) |
| Figure S8 | 24-well | eGFP-p50 | NF-κB-Firefly luciferase (180 ng) + pCMV-Renilla luciferase (60 ng) + eGFP-p50 (30 ng) |
| | | eGFP-p50 $Cry2_{olig}$-mCh-$FUS_N$-NbGFP + $Cry2_{olig}$-mCh-$FUS_N$-NLS | NF-κB-Firefly luciferase (180 ng) + pCMV-Renilla luciferase (60 ng) + eGFP-p50 (30 ng) + pAF301 (180 ng) + pAF082 (180 ng) |
